# NF-κB mediates transactivation of HNRNPD, resulting in PTEN destabilization and constitutive activation of the PI3K-AKT pathway in oral cancer cells

**DOI:** 10.1101/2025.01.05.631360

**Authors:** Vikas Kumar, Anjani Mittal, Anurag Kumar, Moien Rasheed Lone, Vandana Yadav, Priya Verma, Dibyabhaba Pradhan, Deepika Mishra, Alok Thakar, Shyam Singh Chauhan

## Abstract

Heterogeneous Ribonucleoprotein D (hnRNPD), an RNA binding protein transcriptionally upregulated by NF-κB transcription factor, is associated with poor outcome of Oral Squamous Cell Carcinoma (OSCC). However, the role of hnRNPD in OSCC remains elusive. This study reveals that hnRNPD positively affects the proliferation, migration, invasion, and survival of OSCC cells. Transcriptome profiling in hnRNPD knockout cells identified significant upregulation of PTEN and inhibition of the PI3K/AKT/mTOR axis. HnRNPD mediates the destabilization of PTEN mRNA by binding to the class II AU-Rich Element (ARE) in 3’UTR of PTEN. The expression of hnRNPD and PTEN are strongly negatively correlated in OSCC tissue specimens, further corroborating hnRNPD-mediated PTEN destabilization. The hnRNPD knockout inhibited autophagy, evident by an accumulation of autophagic vesicles and decreased autophagic flux. Mechanistically, the hnRNPD knockout reduced the expression of NF-κB, eventually downregulating its transcriptional target LC3b, a key mediator of autophagy. SA-β-Galactosidase staining in hnRNPD KO cells conclusively demonstrated the onset of cellular senescence. The present study demonstrates hnRNPD-driven positive modulation of autophagy via NF-κB, independent of the PI3K/AKT/mTOR axis, highlighting it as a novel therapeutic target for treating oral cancer.

**Graphical Abstract:** 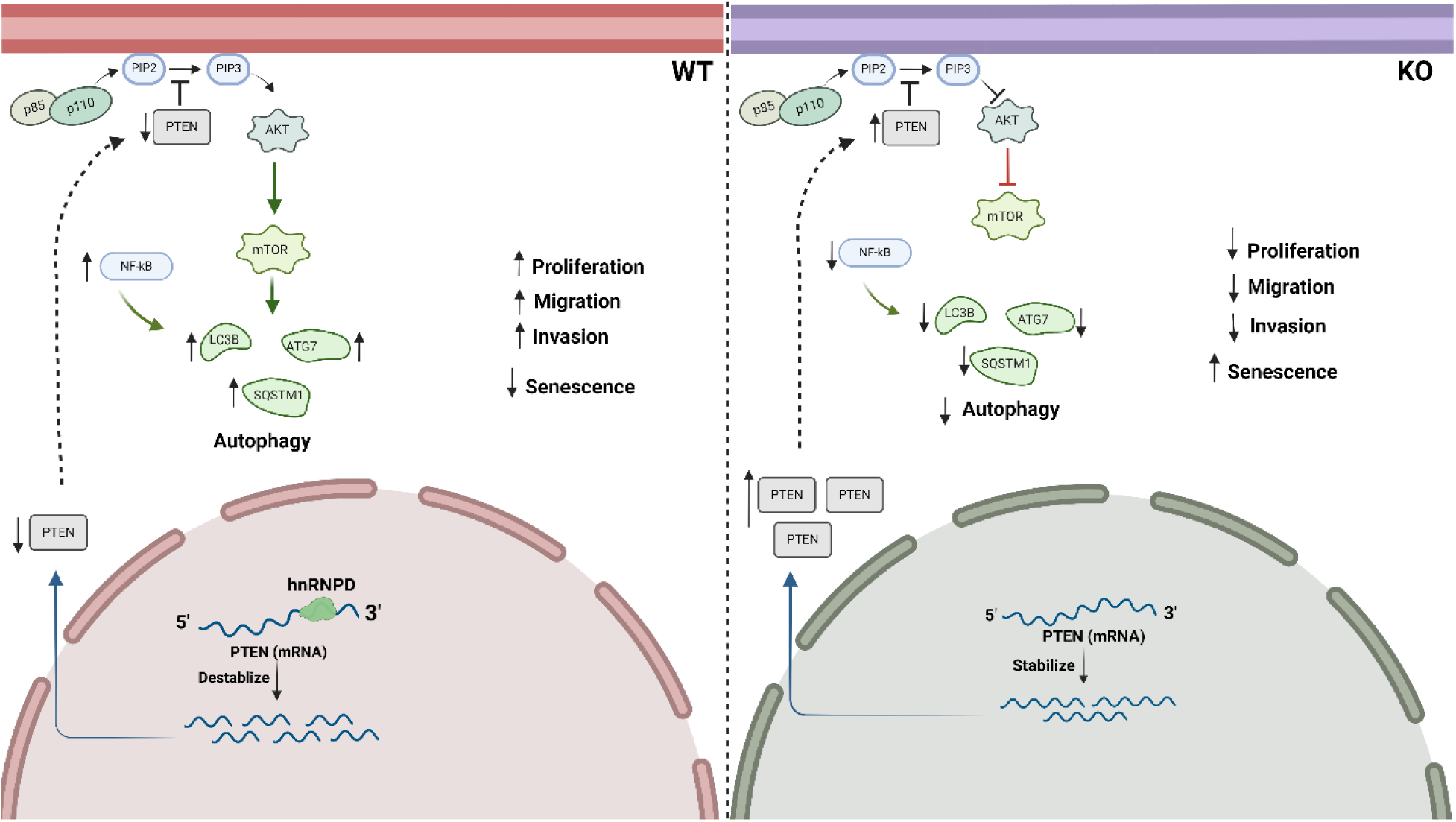

**Highlights:**

- HnRNPD mediates oral cancer cell proliferation, migration, invasion, and survival.
- HnRNPD acts as a novel regulator of the PI3K/AKT/mTOR axis by destabilization of PTEN.
- NF-κB/RelA downregulated on knockout of HNRNPD, inhibiting autophagy through downregulating its transcriptional target LCB-II.
- HnRNPD mediates cellular senescence in oral cancer cells.

## Introduction

The development of Oral Squamous Cell Carcinoma (OSCC) is a complex multistep process involving the alteration of proto-oncogene and tumor suppressor genes^1^. Various molecular signaling pathways, including PI3K-AKT-mTOR, NF-κB, and Wnt, are invariably altered in this malignancy^2,3^. Interestingly, all three of these pathways are also involved in the modulation of autophagy^4–6^. The NF-κB mediates the cross-talk between the PI3K-AKT-mTOR axis by directly regulating the transcription of mTOR^7^. The constitutively active AKT can directly activate its downstream target mTOR that can inhibit autophagy ^8,9^. This is due to the down-regulation of a tumor suppressor gene PTEN, a negative regulator of this cell signaling pathway^10,11^. NF-κB is a key transcription factor that controls the transcription of various genes involved in cellular proliferation, survival, tumorigenesis, and immune and inflammatory responses^12,13^. Increased nuclear expression of NF-κB is associated with metastasis, which accounts for poor survival of oral cancer patients^14^. NF-κB has been reported to mediate the transcription of BECN1, LC3, ATG5, and mTOR, activating autophagy^7^. Downregulation of basal levels of autophagy by changes in the expression of molecular markers can both promote and suppress the survival of oral cancer cells, acting like a dual-edged sword^15–17^. In oral cancer, activated autophagy is associated with chemoresistance via degradation of chemotherapeutic agents^18^. Conversely, autophagic activation can also result in radiosensitivity^19^.

Heterogeneous Nuclear Ribonucleoprotein D (hnRNPD; also known as Adenylate Uridylate rich RNA binding Factor 1 (AUF1), the first identified RNA binding protein, is an ARE (AU-Rich Element) binding protein that regulates the stability of mRNAs encoded by genes involved in tumorigenesis, cell cycle regulation, senescence, cell proliferation, and apoptosis^20–23^. Previous studies report that siRNA-mediated knockdown of hnRNPD decreases cell proliferation, migration, and invasive properties of cancer cells of various origins, including thyroid, colon, skin, esophagus, stomach, breast, bladder, and liver^20–24^. In thyroid carcinoma cells, the knockdown of autophagic protein Beclin-1 increased hnRNPD expression, stabilizing transcription factor ZEB1, resulting in epithelial to mesenchymal transition and promoting invasion^28^. The posttranslational modification of another RNA binding protein, FUS, in glioma cells resulted in hnRNPD-mediated stabilization of autophagic inhibitor ATGD4, thereby inhibiting autophagy^29^. Thus, several reports have demonstrated the involvement of hnRNPD in various cellular processes that promote tumorigenesis and survival of cancer cells^26,27,30–32^. Previously, we have reported the over-expression of hnRNPD in OSCC and established its association with poor prognosis^33^. We have also demonstrated the role of NF-κB in the transcriptional upregulation of hnRNPD in oral cancer cells^34^. However, to date, no systematic study has been carried out to elucidate the role of hnRNPD in oral cancer cells. Therefore, we aimed to elucidate the role of human hnRNPD in oral cancer by employing CRISPR-Cas9 and shRNA-based strategies to knockout (KO) and knockdown (KD) *HNRNPD*, respectively. The work presented here describes the findings of these KO/KD studies and identifies the mediatory role of PTEN, a negative regulator of the PI3K/AKT/mTOR axis, on autophagic flux. Further, we identify a specific role for NF-κB in the transcriptional regulation of autophagy pathway components, concluding that in oral cancers, hnRNPD downregulates NF-κB expression and dysregulates autophagic flux.

## Materials and methods

### Cell culture

Oral squamous cell carcinoma (OSCC) cell line SCC-4 was obtained from American Type Culture Collection (ATCC, VA, USA) and characterized by STR profiling. OSCC cells were grown in monolayer cultures in Dulbecco’s modified eagle medium (DMEM) (Gibco, CA, USA) supplemented with 10 % fetal bovine serum (FBS) (Gibco, CA, USA), 1 mM l-glutamine, 100 μg/ml streptomycin and 100 U/ml penicillin in a humidified CO_2_ incubator (5 % carbon-dioxide, 95 % air) at 37 °C.

### Antibodies

List of antibodies used in the present study and their dilutions were given in supplementary table 1.

### Primers

Custom-synthesized oligonucleotides were purchased from Integrated DNA Technologies (IDT). List of primers used in the present study were given in supplementary table 2.

### Design and cloning of gRNAs into pDG459 vector

Guide RNA for hnRNPD were designed using the CRISPick Genetic Perturbation Portal (https://portals.broadinstitute.org/gppx/crispick/public) an online available software. Guide RNA with the highest score and least number of off-targets was synthesized and used in the present study. Custom-synthesized oligonucleotides for forward and reverse guide RNA were diluted to a final concentration of 100 µM. Then 1 µL 10x T4 ligase buffer, 7 µL ddH2O, 1 µL forward guide oligonucleotide (100 µM) 1 µL reverse guide nucleotide (100 µM) were mixed and heated at 95°C for 5 min and then allowed to cool down to 25°C at the rate of 5^0^C/minute using a PCR machine. Then the annealing mixture was diluted 1:200 with nuclease-free water. Single-step Digestion and Ligation was used to clone annealed oligonucleotides into pDG459vector. For this 100ng of pDG459 vector stock, 1X FastDigest buffer, 10 units of BbsI, 0.5 µL T7 DNA Ligase, 2 µL diluted oligo complex (1:200) in a final volume of 20 µL were subjected to 12 cycles of PCR using the cycling parameter 37°C for 5 min, 21°C for 5 min. The resulting PCR products were used for the transformation and identification of recombinant plasmids.

### HnRNPD knockout in oral cancer cell line by CRISPR/Cas9

Oral cancer cells SCC4 was transfected with pDG459 plasmid containing double gRNA targeting exon1 and exon2 cloned under the U6 promoter. The transfected cells were grown in puromycin-containing media (2.0µg/mL) for 72 hours. Resistant cells were collected and expanded as single-cell colonies in 96-well plate. Then the expression of hnRNPD in 3 independent clones was assessed by western blotting. Simultaneously the expression of β-actin was also assessed by western blotting and used for normalization for equal loading.

### MTS based Cell Proliferation Assay

Cell proliferation was assessed by 3-(4,5-dimethylthiazol-2-yl)-5-(3-carboxymethoxyphenyl)-2-(4-sulfophenyl)-2H-tetrazolium, inner salt; MTS assay. By using CellTiter 96 Aqueous One Solution Cell Proliferation Assay kit (Promega, Madison, WI, USA). For this, 5×10^3^ cells were seeded in each well of 96-well plates and incubated in a CO_2_ incubator. Then at various time intervals (0, 24, 48, and 72 hours) cells were taken out. Absorbance (O.D.) at 490nm was recorded using a micro plate reader.

The formula for calculation of percentage cell proliferation:

Percentage of cell viability = (A_treatment_ − A_blank_)/(A_control_ − A_blank_) × 100% (where, A = absorbance).

### Matrigel Invasion Assay

The ability of tumor cells to invade is one of the hallmarks of the metastatic phenotype. In the present study, Matrigel invasion assay was performed using a reconstituted basement membrane matrix. For this, cells were seeded in serum-free DMEM at a density of 5×10^4^ on a Matrigel (1mg/ml) coated insert of 24 well transwell plate. Each lower chamber was filled with 500 μL medium of 20% FBS which acted as a chemoattractant. After 24 hrs of incubation, the filters were removed, and the non-migrating cells on the upper sides of the filters were removed using a cotton swab. Then the cells on the filter were fixed with 4% paraformaldehyde for 15 min. The cells crossed the matrigel insert and reached the lower side of filters were stained with 0.2% crystal violet for 1 min and counted in five randomly chosen fields (×100 magnification) per well.

### Transwell migration Assay

For this, cells were seeded in a serum-free DMEM at a density of 5×10^4^ in triplicates on a insert of 24 well transwell plate. Each lower chamber was filled with 500 μL medium of 20% FBS which acted as a chemoattractant. After 24 hrs of incubation, the filters were removed, and the non-migrating cells on the upper sides of the filters were detached using cotton swabs. The filters were fixed with 4% paraformaldehyde for 15 min. The cells located in the lower filters were stained with 0.2% crystal violet for 1 min. Invaded cells were counted in five randomly chosen fields (×100 magnification) per well.

### Colony formation assay

Total 10^3^ of SCC-4, SCC4-Control, and SCC-4 hnRNPD KO1/2/3 cells were seeded in six well plate and incubated for 10 days. At the end of incubation these cells are washed wit 1XPBS and stained wit 0.2% crystal violet and counted manually across multiple fields in light microscope.

### Annexin-V/PI Assay

The frequency of apoptosis and necrosis in cells after knockout of hnRNPD was assessed by annexin-V/PI binding assay. Briefly, 8 X 10^4^ SCC4, SCC4-Control and SCC4 hnRNPD KO1/2/3 cells per well were plated in six well plates. Next day cells cycle in these cells was synchronized by keeping in 1% FBS containing DMEM cells incubated for 24 hours. At the end of the incubation, cells were washed followed by stained with annexin-V/PI apoptosis detection kit according to the manufacturer’s instructions. Briefly, the cells were washed twice with PBS, and resuspended in annexin-V binding buffer. To the cell suspension, 5 µl of FITC-conjugated annexin-V and 10 µl of PI (50 µg/ml) solution were added and further incubated for 15 min at room temperature. About 10,000 events were acquired in FACS LSRII flow cytometer. The frequency of annexin-positive and double positive (apoptotic) cells was determined using BD FACS Diva software.

### Western Immunoblotting

Cells were lysed in RIPA buffer (50mM Tris HCl, pH7.5; 1mM EDTA, pH 8.0; 1% Nonidet P-40; 150mM NaCl; 10mM MgCl_2_) containing 1X protease inhibitor cocktail (Sigma Aldrich, St. Louis, MO, USA) and the lysates were centrifuged at 10,000 X g for 10 mins at 4°C to remove the debris. The supernatants were transferred to fresh micro-centrifuge tubes and total amount of protein was estimated by BCA method.Lysates containing equal amount of total protein (∼50 µg) were resolved on 12% SDS-PAGE. The proteins were transferred on to a 0.45µm (pore size) PVDF membrane (mdi, Amballa Cantt, India). After transfer, the membrane was incubated in blocking buffer (5% Bovine serum albumin or 5% non-fat milk dissolved in TBS I) for 2 hour at room temperature. Membrane was followed by incubation with diluted primary antibody in TBS I, containing 1% BSA or 1% NFM overnight at 4°C. Post incubation the membrane was washed (with vigorous agitation) three times (5mins each) with TBS II containing 0.05% tween, followed by TBS II containing 0.02% tween and finally with TBS II alone. The HRP conjugated secondary antibody was then diluted (1:2000) in TBS I containing 1%BSA or 1% NFM. The membrane was then incubated in secondary antibody for 2hrs. at room temperature followed by 3 washing with TBS II tween (0.05%), TBS II tween (0.02%) as described above. The membrane was then covered with ECL solution (Thermo Fisher Scientific Inc. Rockford, IL USA) for 1-5 minutes at room temperature and subjected it to autoradiography for various time periods.

### mRNA Half-life assay

2 x 10^5^ SCC-4 or SCC-4 hnRNPD KO1 cells were seeded per well in 6 well plate. After 24 hours, Actinomycin D at final concentration of 5μg/mL in 10% DMEM was added to each well. Cells were harvested at 0, 2, 4, 6 and 8 hours after the Actinomycin D treatment and processed by RNA isolation using Ribozol as described earlier. An aliquot containing, 500ng of total cellular RNA was reverse transcribed and subjected to qRT-PCR analysis. 18S rRNA was used as the internal control for normalization of Ct values. % mRNA remaining was calculated using 2 –ΔΔCt method. Half-life was calculated using one phase decay equation of Graphpad Prism 6 and p value was calculated using paired two tailed student t test.

### Real-Time PCR

Two μg of total RNA was reverse transcribed using M-MuLV RT (MBI Fermentas, Vilnius, Lithuania) and random hexamers as described earlier. An aliquot containing 200 ng of the total cDNA was subjected to PCR using primers specific for hnRNPD, and PTEN on a Bio-Rad I-cycler (Bio-Rad, Hercules, CA). PCR reactions were carried out in 20 μl total volume containing 1X Maxima SYBR Green qPCR Master Mix (Thermo Scientific, Waltham, USA). PCR conditions comprised 40 cycles of denaturation 94°C for 30 s, annealing at 59°C for 45 s, extension at 72°C for 1 min, and fluorescence recording at 80°C for 30 s. Similarly, 18S cDNA was also amplified using primers specific for 18S which served as the internal control. Melting curve analysis confirmed no primer dimer formation for human hnRNPD or PTEN or 18S under the above-mentioned conditions. The amplification of the expected sizes of PCR products was confirmed by running on agarose gel (1.5%). Cycle threshold (Ct) values were calculated for each PCR and relative fold change was calculated using 2^−ΔΔCt^ method. The relative abundance of the target genes was calculated using the following formula:

Relative RNA = (1+ E)^-ΔΔCt^; where E is the PCR efficiency. For simplicity PCR efficiencies were assumed to be 100%; therefore, this assumption introduced minimal error in the calculation.

### RNA immunoprecipitation (RIP) assays

RIP was conducted with the Magna RIP RNA-Binding Protein Immunoprecipitation Kit (17–700, Millipore) according to the manufacturer’s instructions. Briefly, magnetic beads coated with 5 μg of specific antibodies against rabbit immunoglobulin G (PP64B, Millipore), or hnRNPD (07-260, Millipore) were incubated with prepared cell lysates overnight at 4 °C. Then, the RNA-protein complexes were washed six times and incubated with proteinase K digestion buffer. RNA was finally extracted by phenol-chloroform RNA extraction methods.

### RNA sequencing and differential gene expression analysis

The SCC4 Control, SCC4 hnRNPD KO1, and SCC4 hnRNPD KO2 clones in triplicates was subjected to transcriptome analysis by using the Illumina platform. Followed by RNA-seq analysis was carried out on raw reads of nine transcriptome profiles. RNA-seq data are available at GEO: GSE26XXXX. The raw sequencing reads were quality-checked by FASTQC. The adapter trimming was performed using Trimmomatic-0.36. Quality reports for the filtered reads are generated using MultiQC. The GRCh38 human reference genome was retrieved from NCBI and indexed with splice site aligner hisat2. Subsequently, raw reads are mapped to the reference genome using hisat2, and quantification of aligned transcripts was carried out using STRINGTIE. The normalized read fragments per kilobase of exon per million reads mapped (FPKM) are used for computing differential gene expression analysis using the LIMMA Bioconductor package. The genes with |fold change| >= 2 and P value <= 0.05 are used as a threshold for reporting differentially expressed genes.

### Gene Ontology and Pathway Enrichment Analysis

The differentially expressed genes (DEGs) observed in SCC4 Control, SCC4 hnRNPD KO1 and SCC4 hnRNPD KO2 groups are enriched for gene ontology terms viz. biological processes, molecular function, and cellular component using ClusterProfiler. Furthermore, the involvement of DEGs in diverse metabolic pathways was determined based on Kyoto encyclopedia for genes and genomes (KEGG) database.

### Tissue specimens

After obtaining approval from the Institute Ethics Committee for Post Graduate Research, All India Institute of Medical Sciences, New Delhi, India (NO.IECPG-162/19.04.2018), previously prepared paraffin-embedded tissue specimens were used in the present study as described in our previous publication. Before sample collection, all participants and/or their legal guardian(s) gave written informed consent, a mandatory requirement from the institute’s ethics committee. All experiments were performed according to guidelines issued by the institutional ethics committee. A total of 37 oral cancer tissue specimens were utilized in the present study. Clinico-pathological parameters of OSCC specimens are presented in Supplementary Table S3.

### Immunohistochemistry

Paraffin-embedded tissue sections were deparaffinized followed by antigen retrieval and quenching of endogenous peroxidase activity with hydrogen peroxide (0.3% v/v). Then the non-specific binding was blocked with 1% bovine serum albumin (BSA). The tissue sections were then incubated with the target-specific antibody for 16 h at 4°C. The primary antibody was detected using the Dako Envision kit (Dako CYTOMATION, Glostrup, Denmark) with diaminobenzidine as the chromogen and counterstained with hematoxylin. The sections were evaluated by light microscopy and scored using a semi-quantitative scoring system for both staining intensity (nuclear/cytoplasmic) and percentage positivity. The tissue sections were scored based on the % of immunostained cells as: 0–10% = 0; >10–30% = 1; >30–50% = 2; >50–70% = 3 and >70–100% = 4. Sections were also scored semi-quantitatively on the basis of staining intensity as negative = 0; mild = 1; moderate = 2; intense = 3. Finally, a total score was obtained by adding the score of percentage positivity and intensity giving a score range from 0 to 7 as described previously.

### Statistical analysis

Statistical comparison between two groups was performed using Student’s *t*-test. The Spearman’s correlation was analyzed by using GraphPad Prism 6 software (Graphpad Software, San Diego CA, USA).

## Results

### HnRNPD positively regulates cell proliferation, migration, invasion, and survival of oral cancer cells

To evaluate the functional role of hnRNPD in oral cancer, we utilized CRISPR/Cas-9 mediated gene editing to generated three stable hnRNPD Knockout (KO) clones of the SCC4 cell line. The three knockout clones were confirmed by PCR screening and validated further using western blot analysis (SF1 and Figure 1A). As expected, all three clones (SCC4 hnRNPD KO1, KO2, and KO3) had no expression of hnRNPD protein compared to transfection reagent-treated (mock) and empty-vector transfected (control) cells. To evaluate the role of hnRNPD in cell proliferation, MTS (3-(4,5-dimethylthiazol-2-yl)-5-(3-carboxymethoxyphenyl)-2-(4-sulfophenyl)-2H-tetrazolium)-based cell proliferation assay was performed every 24 hours for 72 hours. A significant reduction (p=0.0134) in cell proliferation of hnRNPD KO clones was observed compared to mock and control cells (Figure 1B). Considering the drastic reduction in cell proliferation, we further evaluated cell migration and invasion using Boyden chamber-based assays. SCC4 hnRNPD KO cells exhibited a drastic reduction in cellular migration (p=0.0006) and invasion (p=0.00078) abilities as compared to control counterparts (Figure 1C-F). Prompted by these findings, we further investigated the colony-forming ability of these cells and found that it was severely compromised in hnRNPD-edited cells compared to control cells (SF2A-B). To evaluate the effect of hnRNPD KO on the processes of apoptosis and necrosis, we performed Annexin V and propidium iodide (PI) staining. The SCC4 mock, SCC4 control, and two independent hnRNPD knockout clones of SCC4 cells were stained with Annexin V/PI and analyzed with flow cytometry. We observed that knockout of hnRNPD led to no significant changes in apoptosis and necrosis (SF2C-D).

**Figure 1.**
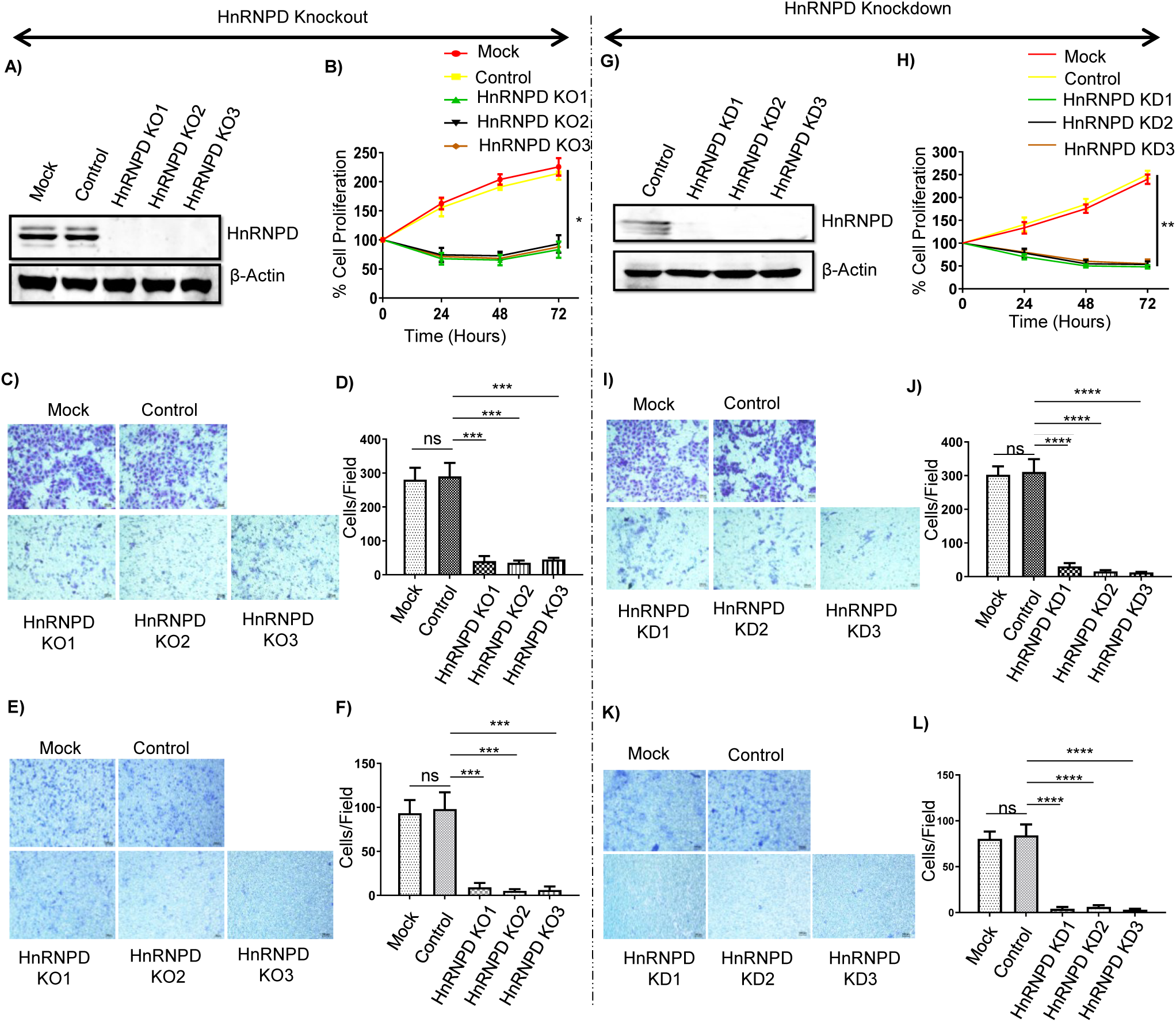
Knockout and knockdown of hnRNPD leads to a reduction in the oncogenic properties of oral cancer cells while no change in apoptosis. **(A)** An equal amount of protein from hnRNPD KO and control (empty vector treated) SCC-4 cells were resolved on 10% SDS-PAGE and subjected to Western blotting using a monoclonal antibody against hnRNPD. Western blot for β-actin served as internal control and was used for normalization for equal loading. **(B)** Effect of hnRNPD deletion on cell proliferation by CellTiter 96® A_Queous_ One Solution Cell Proliferation Assay results of three independent experiments representative as mean±SD in triplicate. **(C, E**) Representative microscopic images of transwell migration and invasion assays followed by staining with 0.2 % crystal violet. (**D, F**) Number of cells was counted in the four different fields and their mean±SD were plotted. (**G**) An equal amount (20µg) of total cell lysate proteins from SCC4 Control, three individual SCC-4 hnRNPD clones were resolved on 10% SDS-PAGE and subjected Western blotting using monoclonal antibodies against hnRNPD. Western blot for β-actin served as internal control and was used for normalization for equal loading. (**H**) Effect of hnRNPD knockdown on cell proliferation by CellTiter 96® A_Queous_ One Solution Cell Proliferation Assay results of three independent experiments representative as mean±SD in triplicate. (**I**, **J**) Representative microscopic images of transwell migration assay and quantification of migratory cells. (**K, L**) Representative microscopic images of transwell invasion assay and quantification of invasive cells. Total numbers of cells were counted in the four different fields and their mean±SD was plotted. All of the results are from three independent experiments performed in triplicates, statistical significance was calculated by unpaired Student’s t-test (*P ≤ 0.05, **P ≤ 0.01, ***P ≤ 0.001, ****P ≤ 0.0001).

To validate the observed phenotypes of reduced cell proliferation, migration, and invasion are true effects of hnRNPD gene editing, we further generated shRNA-mediated stable hnRNPD knockdown (KD) SCC4 cells (Figure 1G). We performed cell proliferation, migration, and invasion assay in these cells and found that knockdown of hnRNPD resulted in a significant reduction of cell proliferation, migration, and invasion (Figure 1H-L). These are similar observations to the hnRNPD KO SCC4 cells, confirming that changes in cellular functions of hnRNPD KO cells are the true effect of hnRNPD gene editing.

### RNA-Sequencing revealed that hnRNPD plays a key role in autophagy through modulation of tumor suppressor PTEN and mTOR signaling pathway

To further elucidate the effect of hnRNPD loss, HnRNPD-KO SCC4 oral cancer cells were subjected to RNA sequencing for transcriptome analysis. Differentially expressed genes (DEGs) in hnRNPD KO cells were identified from 240434 transcripts captured. A total of 3778 transcripts were found to be dysregulated in hnRNPD KO cells (Figure 2A). Transcriptome analysis revealed that out of the 3778 DEGs, 1567 are upregulated while the remaining 2211 genes are down-regulated. 3439 DEGs were found to be common in both hnRNPD KO clones, while 319 and 235 unique DEGs were detected in SCC4 hnRNPD KO1 and 2 respectively (Figure 2B). Hierarchical clustering analysis on DEGs revealed that these samples belong to two groups: SCC-4 Control and SCC-4 hnRNPD KO clones (Figure 2C). Among these DEGs, we detected some genes (PTEN, YWHAZ, BECN1, and SQSTM1) that play key roles in autophagy (Table 1). KEGG gene ontology analysis was performed to identify the functional profiles of DEGs; this analysis revealed that decreased hnRNPD expression is highly associated with autophagy and the mTOR pathway (Figure 2D). This suggests that hnRNPD plays a major role in the regulation of autophagy.

**Figure 2.**
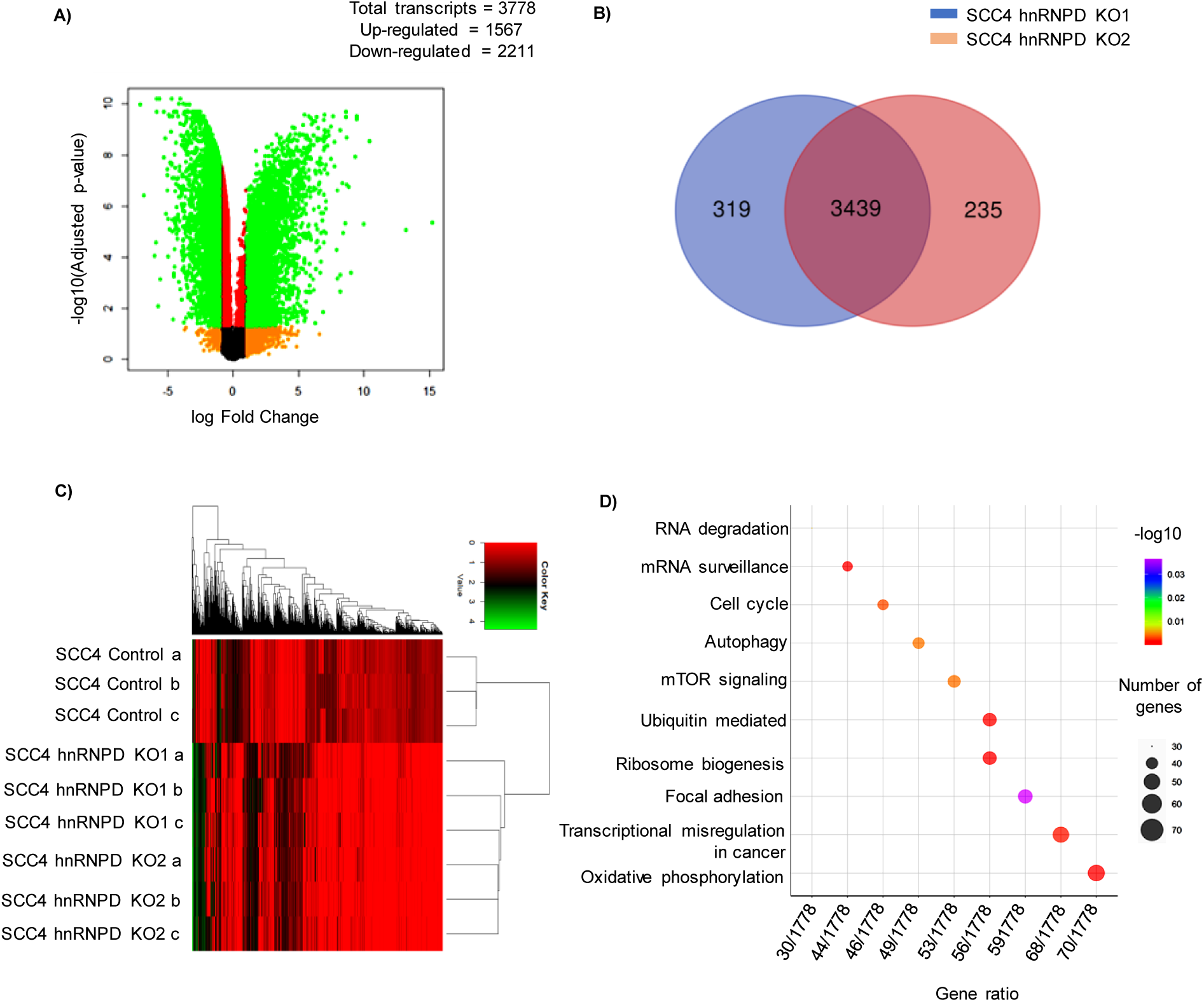
RNA-sequencing suggests the role of hnRNPD in autophagy. **(A)** Volcano plots showing the deferentially expressed genes (DEGs) in SCC-4 hnRNPD knock out clones (KO1 & KO2) in comparison of SCC-4 control cells. This graph is plotted negative logarithm of false discovery rate on y-axis, while log fold change onto x-axis. Right side of this graph represents up-regulated genes, whereas on the left side it shows down-regulated genes. Data represented here is from RNA-Seq data of N=3 performed in triplicates. **(B)** Venn diagram representing the number of DEGs between SCC-4 hnRNPD KO1 and SCC-4 hnRNPD KO2. **(C)** Heatmap of hierarchical clustering indicates DEGs between SCC-4 Control, SCC-4 hnRNPD KO1 and SCC-4 hnRNPD KO2 (Fold-change > 2, P < 0.05). **(D)** KEGG pathway analysis was performed in total (up-regulated and down-regulated) DEGs. This analysis demonstrates the enrichment of genes involved in autophagy, mTOR, cell cycle and RNA degradation pathways.

**Table 1.**
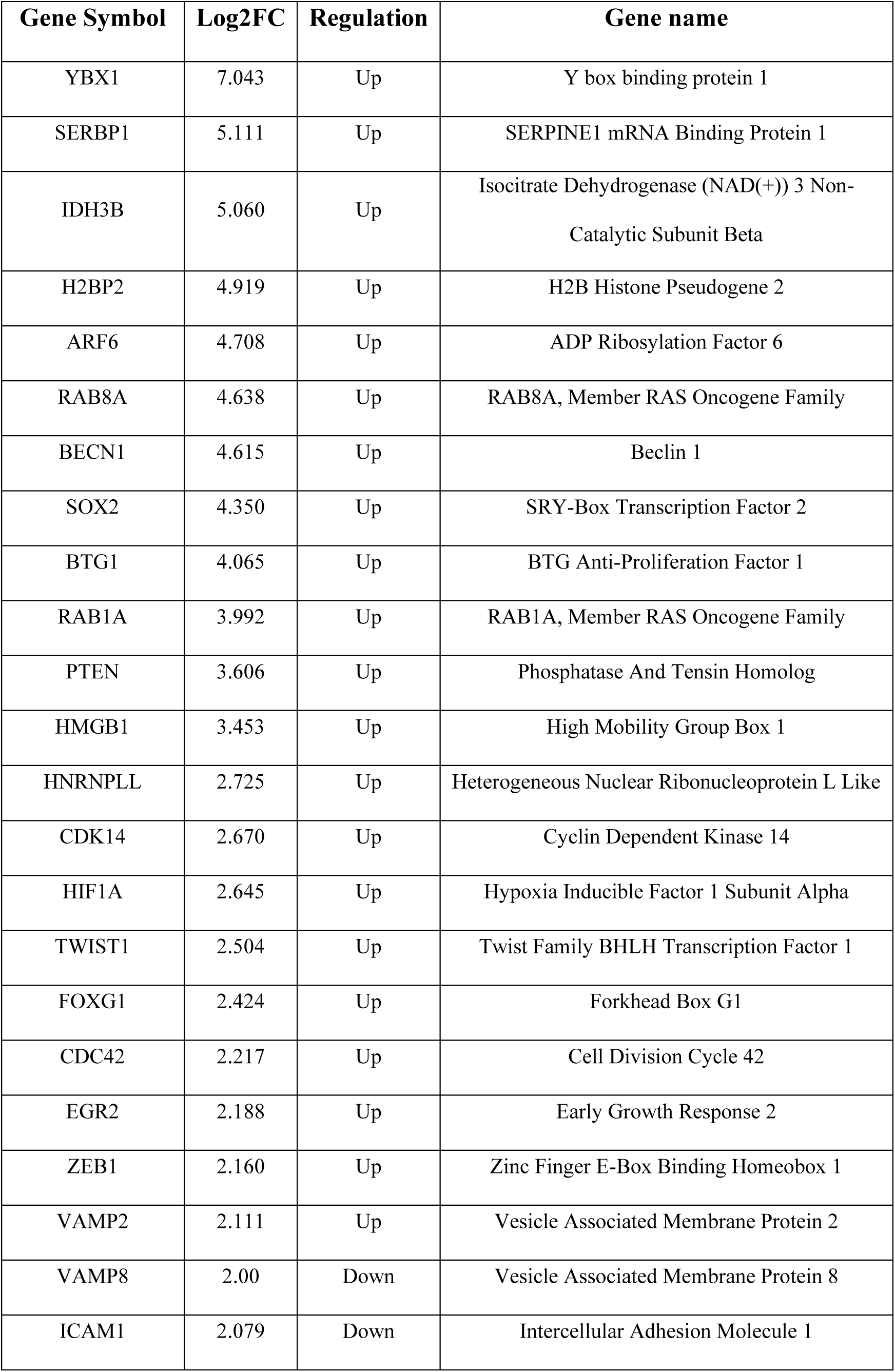
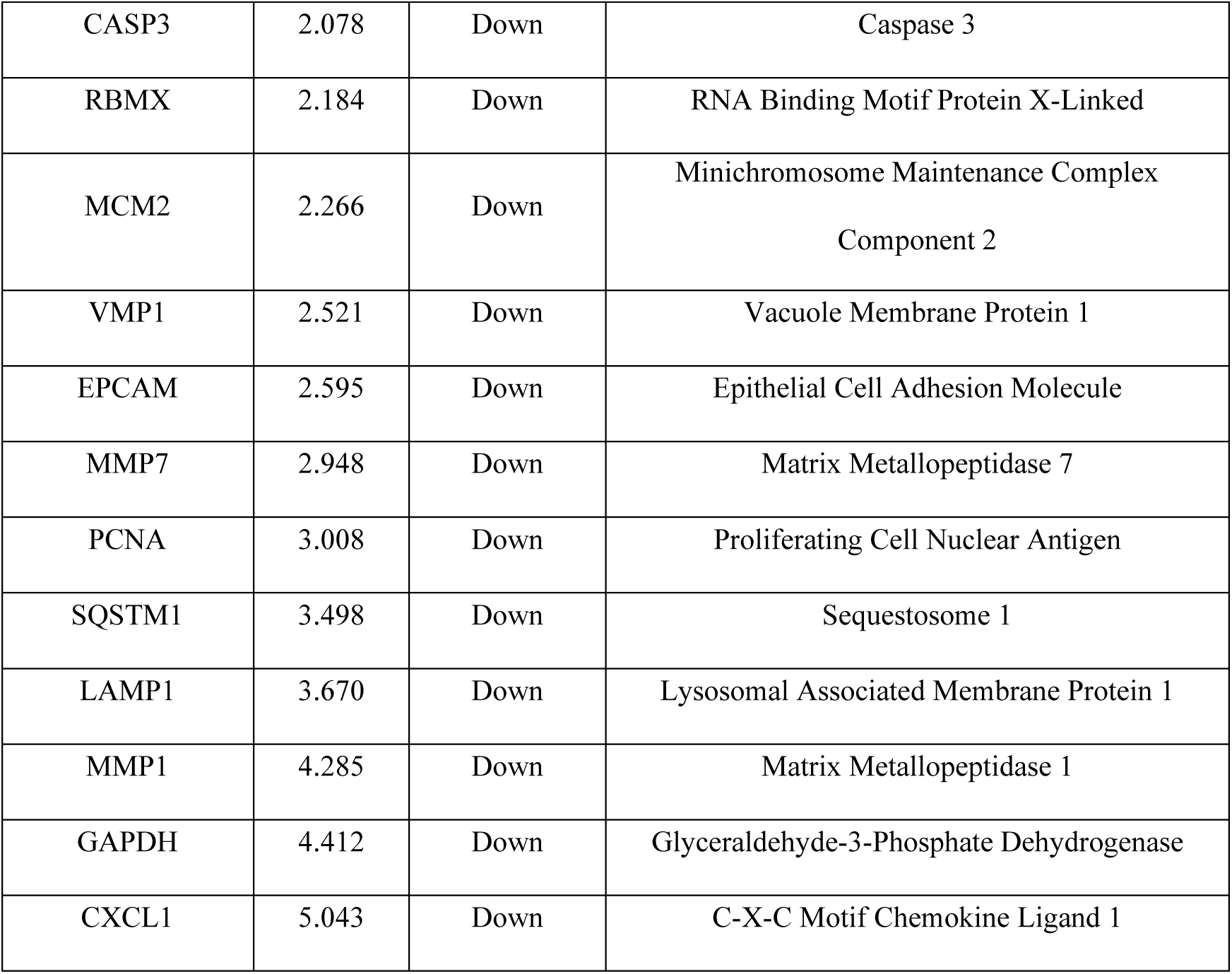
List of significantly dysregulated genes in hnRNPD knockout oral cancer cells.

### HnRNPD knockout led to inhibition of PI3K/AKT/mTOR axis by upregulation of PTEN expression

Knockout of hnRNPD in oral cancer cells led to a decrease in cancerous properties and dysregulation of genes involved in autophagy as revealed by transcriptome analysis. Alterations in this PI3K/AKT/mTOR pathway influence the process of autophagy^17,35^. We next wanted to validate these findings by western blot analysis of the PI3K/AKT/mTOR pathway. Knockout of hnRNPD led to an increase in PTEN expression, which is known to negatively regulate PI3K/AKT/mTOR axis by dephosphorylation of PIP3 to PIP2 (Figure 3). In contrast, PI3K serves as a positive regulator of PI3K/AKT/mTOR axis, which catalyzes the phosphorylation of PIP2 to PIP3 by its catalytic subunit PI3KCA^17^. Specifically, knockout of hnRNPD led to a reduction in the PI3KCA expression and inhibition of this axis (Figure 3A-B). Simultaneously, we observed a decrease in phosphorylation of AKT at serine at position 473 (Phos-AKT Ser473), the fully activated form of AKT; however, no change in the total AKT was observed. Furthermore, a decrease in the mTOR phosphorylation at the serine 2448 position (Phos-mTOR Ser2448) was also observed by using a monoclonal antibody raised against mTOR phosphorylated at S2448. In addition, a decrease in AKT also results in the activation of total GSK3β. Due to a decrease in the inhibitory phosphorylation of GSK3β at serine 9 residues (Phos-GSK3β Ser9). These observations demonstrate that hnRNPD is a positive regulator of the PI3K/AKT/mTOR axis in oral cancer cells.

**Figure 3.**
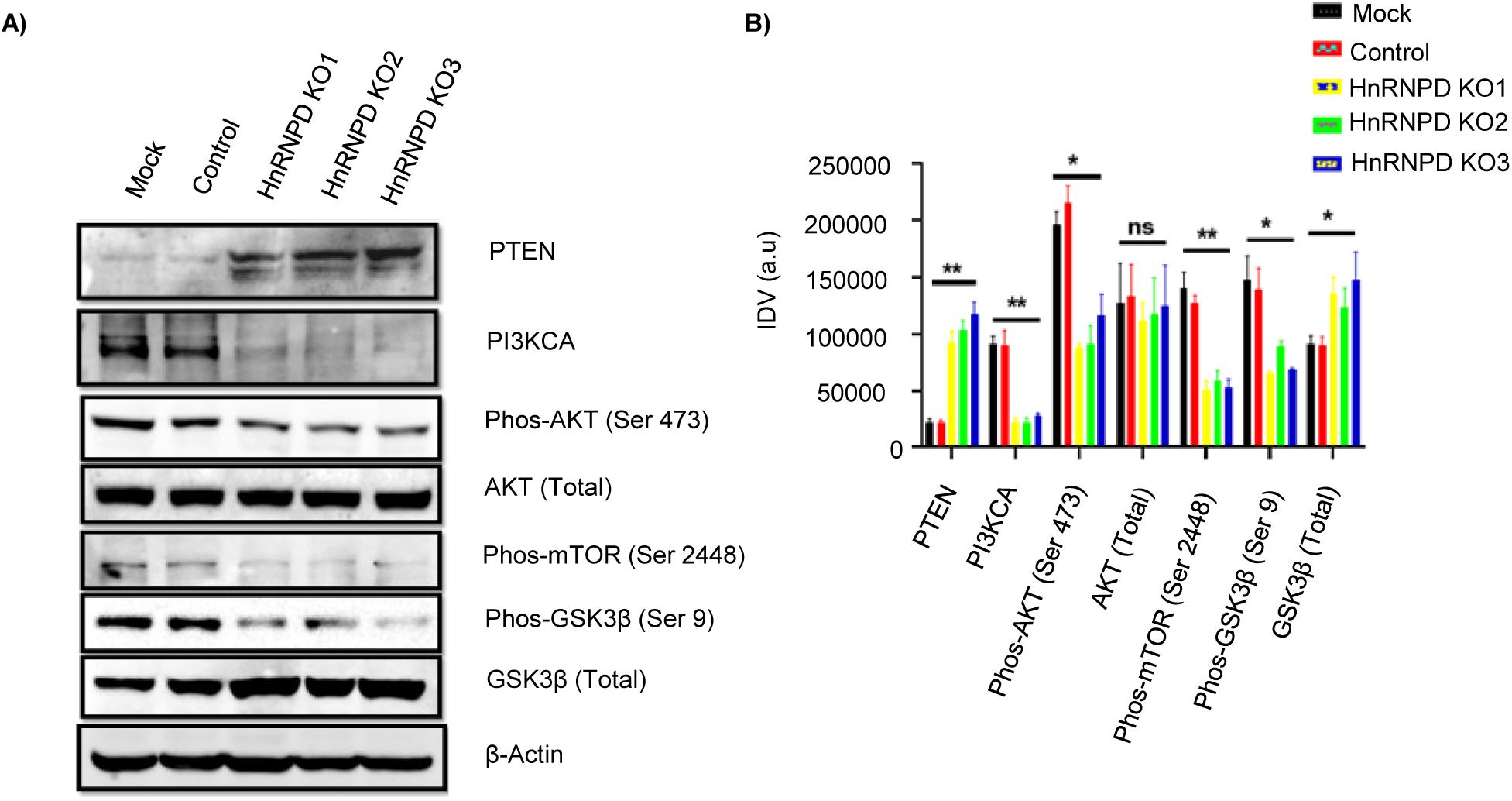
CRISPR/Cas9 mediated knockout of hnRNPD resulted in the inhibition of PI3K/AKT/mTOR axis. 10^9^ SCC-4, SCC-4 control (pDG459), and three independent hnRNPD knockout clones (KO1, KO2, and KO3) cells were seeded in T75 cell culture flasks and placed in a Co2 incubator. After 24 hours the cells were washed with ice-cold Phosphate Buffered Saline, pH-7.4 (PBS), followed by cell lysis in RIPA buffer, and cell lysates were cleared of cell debris by centrifugation at 3000Xg. An equal amount (20µg) of protein from these cells was resolved on 10% SDS-PAGE and subjected to Western blotting using specific antibodies for each protein. Specific bands were quantitated densitometrically using Image J software (NIH, U.S.A). The β-actin served as internal control and was used for normalization for equal loading. **(A)** PI3K/AKT/mTOR axis. (B) Densitometric quantification of PI3K/AKT/mTOR axis. All of the results are from three independent experiments performed in triplicates, statistical significance was calculated by unpaired Student’s t-test (*P ≤ 0.05, **P ≤ 0.01, ***P ≤ 0.001, ****P ≤ 0.0001).

### HnRNPD-mediates PTEN mRNA de-stabilization in oral cancer cells

The RNA-Seq analysis in hnRNPD knockout cells revealed a 3.6-fold increase in the levels of PTEN mRNA (Table1). The knockout of hnRNPD led to an increase in PTEN protein expression, which resulted in the inhibition of PI3K/AKT/mTOR axis (Figure 3A). These results were further corroborated by measuring transcript levels of PTEN in these cells; a significant ∼12-fold increase (p=0.00005) in the PTEN mRNA levels was measured in comparison to SCC-4 mock and SCC-4 control cells (Figure 4A). As hnRNPD is an RNA binding protein well known to regulate mRNA stability, we sought to investigate its role in regulating PTEN mRNA stability. We utilized the publically available POSTAR3, a CLIPdb database (http://111.198.139.65/RBP.html) for the identification of the hnRNPD binding motif in the PTEN mRNA 3’UTR. This analysis revealed the presence of three overlapping conserved pentameric (AUUUA) hnRNPD binding motifs that belong to class II AU-Rich Element (ARE) in the 3’UTR of PTEN mRNA (Figure 4B). Class II AREs are involved in mRNA destabilization. We measured the half-life of PTEN mRNA in hnRNPD knockout cells and found the half-life of PTEN mRNA in SCC-4 control cells was observed to be 3.53 hours, whereas in SCC-4 hnRNPD knockout cells half-life increased to 5.47 hours (Figure 4C). These results convincingly demonstrate hnRNPD mediated destabilization of PTEN mRNA.

**Figure 4.**
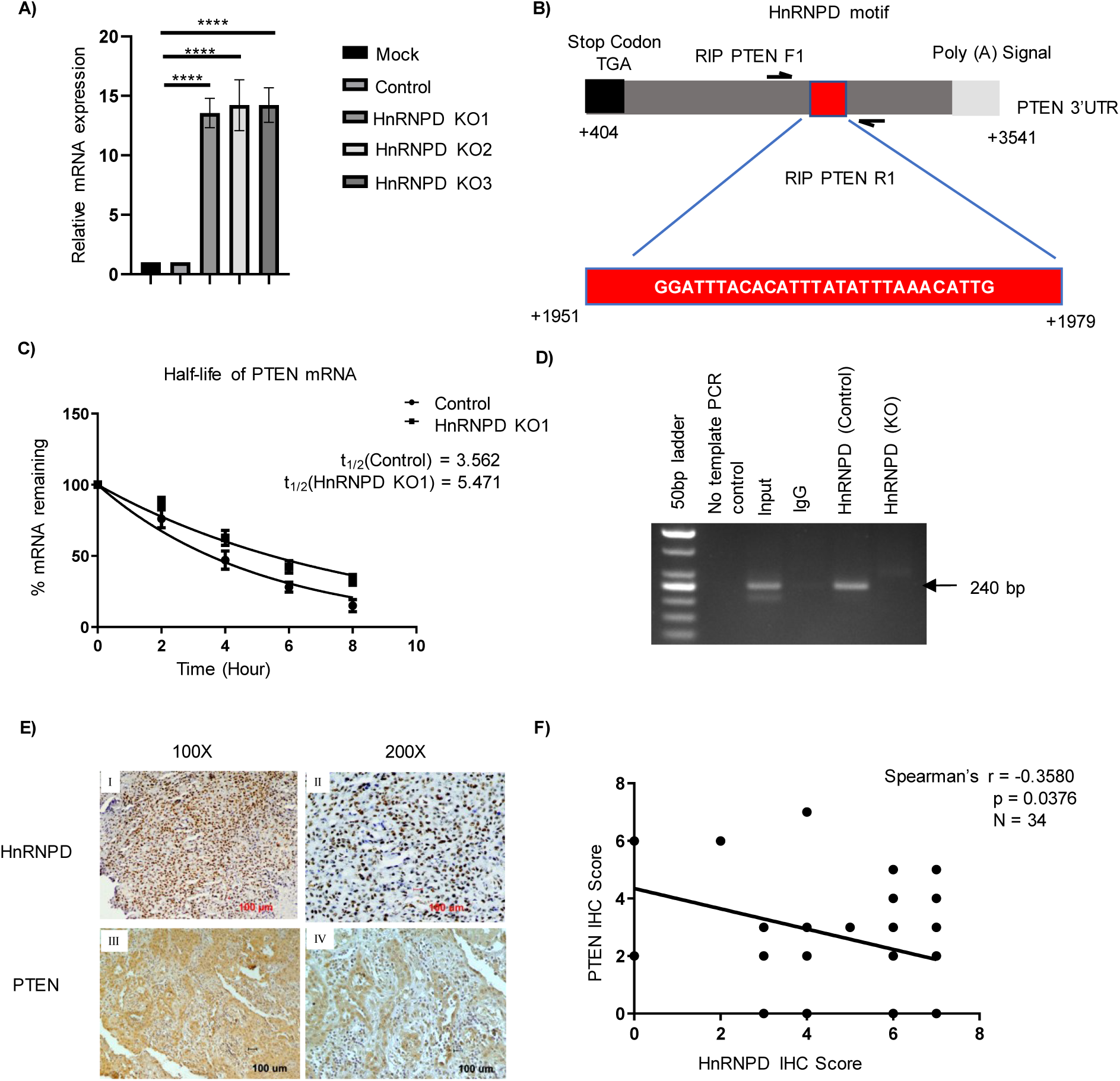
HnRNPD mediates destabilization of PTEN mRNA in oral cancer cells. **(A)** Total RNA isolated from SCC-4 cells and SCC-4 hnRNPD KO clones was reverse transcribed and subjected to real time PCR using specific PTEN primers. Simultaneously, the PCR was also performed using 18S ribosomal RNA specific primers and served as internal control for normalization of PTEN transcript. **(B)** Schematic representation depicting the positions of hnRNPD binding motif in the PTEN 3’UTR and primers used for RIP assay. **(C)** The SCC-4 control and SCC-4 hnRNPD KO1 cells were treated with 5μg/ml of actinomycin D for various time points to block mRNA synthesis. Total RNA was harvested followed by the qPCR as described in the text. Half-life was calculated using non-linear one phase exponential decay equation inGraphpad Prism 6.0. **(D)** HnRNPD bound mRNA was immunoprecipitated from SCC-4 control and SCC-4 hnRNPD KO cells with anti-hnRNPD antibody and subjected PCR after elution of bound mRNA from hnRNPD using the primers depicted in the figure 24C. Total cellular RNA isolated from SCC-4 cells was used as input. PCR with gene specific primers and RNA immunoprecipitated using normal rabbit IgG served as negative control (IgG lane). Similarly, PCR performed with gene specific primers without a template also served as control. A 50 bp DNA ladder was used to determine the size of the amplified PCR fragments on agarose gel. Representative agarose gel image of RT-PCR fragment using RIP assay for hnRNPD binding motif on PTEN mRNA. Values are mean ± SD from three independent experiments. Values significantly different from respective controls have been marked by stars (**P ≤ 0.01, ****P ≤ 0.0001). Expression of PTEN and hnRNPD in oral cancer. **(E)** Paraffin-embedded tissue sections of oral cancer were immunostained using anti-hnRNPD or anti-PTEN antibody. The expressions of both the proteins were scored independently by two pathologists blinded to the identity of sections and their score. The representative images of OSCC immunostained sections. Nuclear expression of hnRNPD (I-II) in OSCC, whereas mild cytoplasmic staining of PTEN (III-IV). **(F)** Correlation between hnRNPD and PTEN expression in OSCC tissue specimens was assessed by Spearmen’s correlation analysis. This analysis revealed a strong negative correlation between nuclear expression of hnRNPD and cytoplasmic expression of PTEN in OSCC. Values are mean ± SD from three independent experiments. Values significantly different from respective controls have been marked by stars (**P ≤ 0.01, ****P ≤ 0.0001).

A RNA Immunoprecipitation (RIP) assay was performed to confirm the binding of hnRNPD to its cognate binding motif on 3’UTR of PTEN mRNA (Figure 4B). For RIP assay, the mRNA-hnRNPD complex was immunoprecipitated using hnRNPD antibody and incubated with proteinase K to remove the RNA-bound proteins antibody complex. The resulting mixture was used for the isolation of immunoprecipitated RNA, which was reverse transcribed and subjected to PCR using PTEN RIP forward and reverse primers. As evident from these results, amplification of the specific 240 bp PTEN transcript in the SCC-4 cells resulted in an intense PCR fragment when cDNA derived from RNA immunoprecipitated and the hnRNPD antibody was used as a template (Figure 4D). Similarly, no amplification was observed with the cDNA template combined with rabbit IgG isotype control antibody. Importantly, no amplification was observed when RNA immunoprecipitation from the SCC-4 hnRNPD KO cells was subjected to PCR using the same set of primers. A lack of amplification of PTEN transcript in these cells further confirmed the specificity of hnRNPD antibody to hnRNPD protein, demonstrating the specific binding of hnRNPD to its cognate motif in the 3’UTR of PTEN mRNA.

In view of these findings on PTEN mRNA destabilization by hnRNPD, it was of interest to validate these results in samples directly from oral cancer patients. Therefore, we analyzed the expression of hnRNPD and PTEN in clinical specimens from 34 oral cancer patients by immunohistochemistry. Clinical features of the patients used in the present study are given in Table 3. Photomicrographs of tissue sections immunostained with PTEN and hnRNPD antibodies from oral cancer tissue specimens displayed elevated expression of hnRNPD in the nuclear compartment, while a low level of PTEN expression was observed in the cytosol (Figure 4E, I-IV). Spearman’s correlation analysis revealed a strong negative correlation (p=0.0376; r= -0.3580) between hnRNPD and PTEN expression in oral cancer tissue (Figure 4F). These results further corroborate hnRNPD mediated destabilization of PTEN mRNA in oral cancer cells.

### Knockout of hnRNPD in oral cancer cells shifts cells from autophagy to senescence

Results presented thus far suggest involvement of autophagy in hnRNPD knockout SCC-4 oral cancer cells due to dysregulation of PTEN and the PI3K/AKT/mTOR pathway. It made sense, therefore, to perform live cell staining of autophagic vacuoles by CYTO-ID green dye in the hnRNPD knockout and control cells and analyze by flow cytometry. Knockout of hnRNPD led to a 2-fold significant increase (p=0.00001) in the number of autophagic vesicles (Figure 5A). This suggests impairment in the process of autophagy. To further validate the effect of hnRNPD knockout in the process of autophagy, we performed an autophagic flux assay using Bafilomycin A1, a specific inhibitor of vacuolar H^+^ ATPase and macrolide antibiotic that has been traditionally used for the inhibition of autophagy. Treatment of SCC-4 control cells with bafilomycin A1 led to a significant increase in LC3-II, a classical marker of autophagy, but no such protein was detected in SCC-4 hnRNPD knockout cells after bafilomycin A1 treatment (Figure 5B-C).

**Figure 5:**
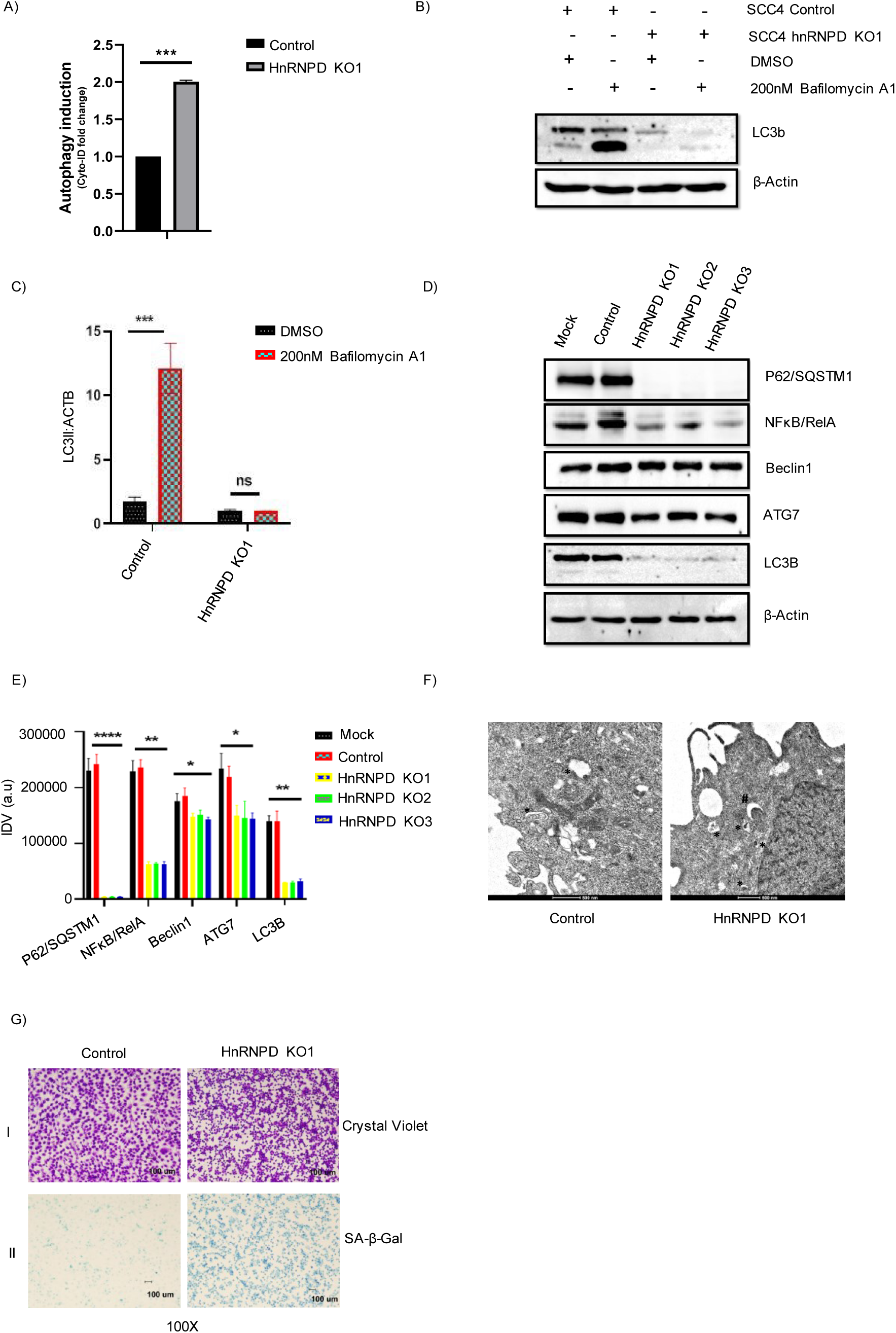
Knockout of hnRNPD results inhibition of autophagy and activation of cellular senescence. Formation of autophagic vacuoles in hnRNPD KO and control SCC-4 cells was evaluated by CYTO-ID quantification kit. For this 10^6^ hnRNPD KO or control cells were incubated for 24 hours in a CO_2_ incubator. The next day, cells were stained with green CYTO-ID for 20 minutes. After incubation cells were processed for flow cytometric analysis or confocal microscopy. **(A)** Quantification of CYTO-ID fold change by flow cytometry. **(B)** HnRNPD KO and control SCC-4 cells treated with bafilomycin A1(200nM) or DMSO followed by Western blotting. **(C)** Densitometric quantification of autophagic flux. **(D)** Equal amount of protein from hnRNPD KO and control (empty vector treated) SCC-4 cells were resolved on 10% SDS-PAGE and subjected to Western blotting to assess the expression of autophagy markers (NF-κB, SQSTM1, Beclin1, ATG7, and LC3b). The β-actin served as internal control and was used for normalization for equal loading. **(E)** Densitometric quantification of autophagic markers **(F)** Representative TEM images of hnRNPD KO and control cells. The ultrastructure of autophagic bodies (astric *) and phagophores (hash #) are marked in the picture (scale bar, 500nm). (G) The Senescence associated β-Galactosidase (SA-β-Gal) staining in SCC-4 control and SCC-4 hnRNPD knockout cells (I). Crystal violet staining (II). All of the results are from three independent experiments performed in triplicates, statistical significance was calculated by unpaired Student’s t-test (*P ≤ 0.05, **P ≤ 0.01, ***P ≤ 0.001, ****P ≤ 0.0001).

Previous reports suggested NF-κB transcription factor mediates the transcriptional upregulation of LC3-II/B, thus promoting autophagy^7^. It was of interest therefore to assess the expression of this transcription factor in addition to autophagy markers in hnRNPD knockout cells. HnRNPD knockout cells displayed a drastic reduction in the expression of NF-κB/RelA (p=0.0069), P62/SQSTM1 (p=0.000006) and LC3B (p=0.006), while Beclin1 and ATG-7 displayed a less dramatic but statistically significant decrease in SCC-4 hnRNPD knockout cells as compared to in SCC-4 mock and control cells (Figure 5D-E. The observed decrease in the NF-κB transcription factor level correlates with a decrease in its transcriptional target LC3B. Furthermore, Transmission Electron Microscopy (TEM) of hnRNPD knockout SCC-4 cells and control cells revealed the presence of phagophore formation (marked by #) and autophagic bodies (marked by *), whereas in control cells only autophagic bodies could be seen (Figure 5F). Conclusively, it demonstrates hnRNPD knockout resulted inhibition of macroautophagy in oral cancer cells by a decrease in the expression of autophagy markers and autophagic flux.

Previously conducted research in the field of aging extensively demonstrates that increased hnRNPD expression is associated with inhibition of cellular senescence^36^. Therefore, it was of interest to evaluate the role of hnRNPD in the cellular senescence of oral cancer cells, as the hnRNPD knockout resulted in the inhibition of autophagy and no change in apoptosis of oral cancer cells. We performed β-Galactosidase staining in SCC-4 control and hnRNPD knockout SCC-4 cells. The hnRNPD knockout SCC-4 cells showed a drastic increase in β-Galactosidase staining as compared to SCC-4 control cells (Figure 5G-I). Simultaneously, we also did a crystal violet staining in an equal number of cells to show the effect of hnRNPD knockout on senescence (Figure 5G-II). The present finding conclusively demonstrates the hnRNPD knockout resulted in the cellular senescence of oral cancer cells. The findings of this study are summarized in Figure 6.

**Figure 6:**
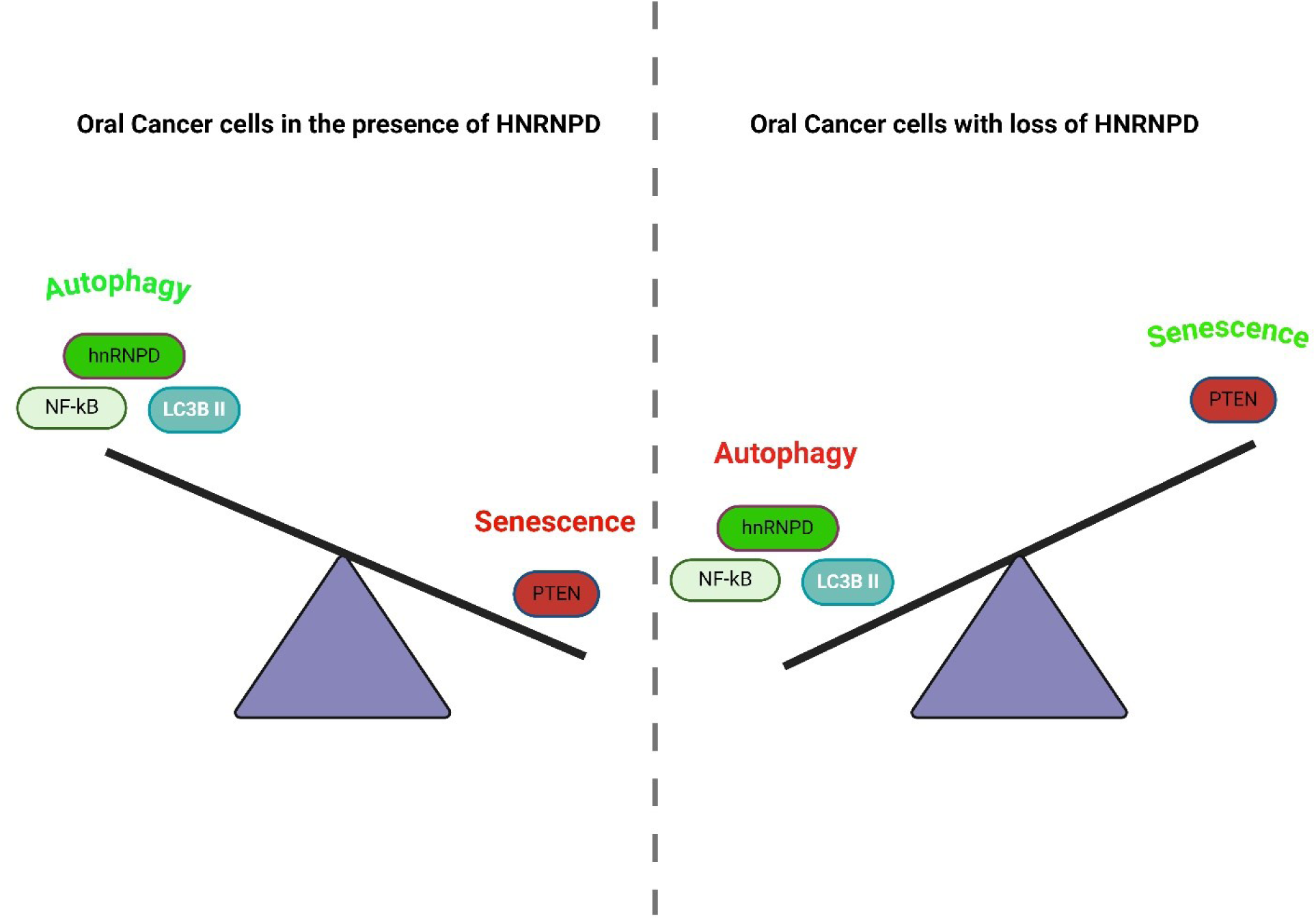
Cell signaling mechanism of oral cancer cells in the presence and absence of hnRNPD. In oral cancer cells, hnRNPD mediates the destabilization of PTEN mRNA by binding to the AU-Rich binding motif, which leads to continuous activation of the PI3K/AKT/mTOR axis that results in constitutive activation of autophagy through NF-kB. In the absence of hnRNPD, the PTEN mRNA stabilized, which led to the inhibition of the PI3K/AKT/mTOR axis and a decrease in the NF-kB levels. This results in the inhibition of autophagy and activation of cellular senescence (Image created with Biorender.com).

## Discussion

HnRNPD plays an essential role in the post-transcription regulation of genes implicated in carcinogenesis. Its over-expression in various malignancies is extensively documented^24,37–41^. We previously reported elevated expression of hnRNPD in OSCC tissue samples and established its association with poor outcomes of the disease^33^. However, very little information about its role in oral cancer was available. Therefore, in the present study, we employed CRISPR/Cas9 technology to elucidate the role of hnRNPD in OSCC-derived SCC-4 cells.

Transcriptome analysis in the hnRNPD knockout SCC-4 oral cancer cells was performed to study the role of hnRNPD in differential gene expression, and pathway enrichment. Previously, various studies have employed a similar strategy to identify the role of hnRNPD in different cancers ^25,32^. This analysis revealed differential expression of 3778 genes after hnRNPD knockout, out of which 1567 were found to be up-regulated while the remaining 2211 were down-regulated. The knockout of hnRNPD led to a significant decrease in the expression of genes that play a crucial role in proliferation (PCNA and MCM2), migration, and invasion (ICAM, EPCAM, and MMP1). Additionally, both upregulation (BECN1 and HMGB1) and downregulation (SQSTM1/P62) expression of the gene involved in the process of autophagy was evident. This analysis further revealed upregulation of the tumor suppressor gene PTEN gene. It is also identified as one of the hub genes in protein-protein interaction network analysis. PTEN is a phosphatase that can dephosphorylate both lipids and proteins ^42–45^. It’s the second highest frequently mutated gene in cancers after p53 ^46,47^. Phosphatase-dependent and independent activities of PTEN govern a majority of cellular processes like maintenance of genome stability, cell proliferation, migration, survival, and metabolism ^43–45^. Immunohistochemical staining and bisulfite sequencing studies in oral cancer revealed that PTEN is more commonly down-regulated rather than mutated at the genomic level^48^. In 30% of OSCC cases, the aberration in the PI3K/AKT pathway leading to enhanced cell proliferation has been observed due to the downregulation of PTEN expression ^49–51^.

Pathway enrichment analysis of differentially expressed genes in hnRNPD knockout cells as compared to control cells revealed the involvement of hnRNPD in biological processes such as RNA splicing, translation initiation, ribonucleoprotein complex biogenesis, and assembly. Transcriptome analysis suggests an additional role of hnRNPD in cellular and molecular processes such as dysregulation of transcription as well as coregulation activity, ubiquitin protein ligase binding, electron transfer activity, transcription factor activity (RNA binding), formation of focal adhesion junctions, and spliceosomal complex, and DNA packaging. Our findings are in agreement with previous reports that hnRNPD mediates stabilization or decay of target mRNA by enhancing the mRNA stability, while it also acts as a transcription factor and mediates the transcriptional upregulation of a target gene ^21,41,52^. Apart from this, it also mediates translational regulation and plays the role of a splicing factor by which it mediates the alternative splicing of mRNA ^53–55^.

Transcriptome analysis in hnRNPD knockout oral cancer cells compared to control cells also revealed its role in the dysregulation of oncogenic properties such as proliferation, migration, and invasion of oral cancer cells. It was of interest to validate these findings in hnRNPD knockout oral cancer cells as compared to control cells, as CRISPR/Cas9 mediated knockout has been reported to have off-target effects ^56^. To rule out the off-target effects in the phenotype observed in CRISPR/Cas9 mediated hnRNPD knockout SCC-4 cells, we also silenced its expression in these cells by shRNA technology. Changes in proliferation, migration, and invasion of oral cancer cells upon shRNA-mediated gene silencing of hnRNPD were similar to those of knockdowns, ruling out the off-target effects of CRISPR/Cas9. Decreased proliferation, migration, and invasion observed in hnRNPD knockout cells agree with a significant decrease in the expression of genes involved in proliferation (PCNA and MCM2), migration (CTTN, RAC1, and ICAM1), and invasion (MMP1, MMP7, MMP10, MMP14, and CTSB) revealed by transcriptome analysis. Our observations are consistent with previous reports showing decreased proliferation, migration, and invasion of thyroid cancer cells upon hnRNPD knockout ^23^. The knockdown of hnRNPD suppressed the proliferation, migration, and invasion of colorectal cancer cells, thereby inhibiting tumorigenesis ^26^. Similar changes after hnRNPD knockdown have also been reported in skin cells and esophagus cancer ^32^. The result of the present study demonstrates a significant decrease in the colony formation ability of SCC-4 after hnRNPD knockout. This observation is in agreement with a substantial reduction in the MCAM and EPCAM expression in transcriptome analysis of hnRNPD knockout oral cancer cells. Consistent with our previous findings, knockdown of hnRNPD has been reported to decrease the colony-forming ability of skin fibroblast, keratinocyte, ESCC, and hepatocellular breast cancer cells ^25,27,32^. Interestingly, the knockout of hnRNPD in oral cancer cells did not have any effects on apoptosis or necrosis. But knockdown of this gene in the thyroid, breast, skin, colorectal, esophagus, ovarian, and hepatocellular carcinoma cells leads to decreased apoptosis (Gao *et al.,* 2016; Tian *et al.,* 2020; X *et al.,* 2021; Zhang *et al.,* 2022) suggesting a context-dependent role of hnRNPD in different cancers.

We observed inhibition of the PI3K/AKT/mTOR axis by upregulation of PTEN expression in hnRNPD knockout cells. PI3K-AKT-mTOR cell signaling pathway has been reported to be constitutively active in oral cancer cells. It is responsible for oral cancer tumorigenesis by promoting oncogenic properties such as proliferation, colony formation, migration, invasion, and inhibiting apoptosis ^3,57–59^. Our results are in agreement with the findings where hnRNPD knockdown has been demonstrated to decrease phosphorylation of AKT at the ser473 position, thereby inhibiting the PI3K/AKT/mTOR axis in cervical cancer cells. Another study reported suppression of tumorigenesis of colorectal cancer cells upon knockdown of hnRNPD via reduced phosphorylation of AKT ^26^.

In the present study, hnRNPD knockout cells displayed significantly higher levels of PTEN mRNA than the control cells due to their increased stability/half-life. HnRNPD is well-known to regulate mRNA stability after binding to its cognate motif on the transcripts. We identified the hnRNPD binding motif in the 3’UTR of PTEN mRNA and established their binding by RIP assay. Therefore, from our results, it can be concluded that hnRNPD post-transcriptionally down-regulates the PTEN expression in oral cancer cells and promotes tumorigenesis. This observation is further corroborated by a negative correlation between hnRNPD and PTEN expression in OSCC clinical tissue specimens. These observations demonstrate a novel mechanism of hnRNPD-mediated regulation of PTEN expression in oral cancer cells.

Previous reports extensively demonstrated that an increase in autophagy is associated with tumor initiation and progression of oral squamous cell carcinoma. Higher expressions of LC3b and p62/SQSTM1 are correlated with poor prognosis of the disease^60–62^. The NF-κB is a key transcription factor that mediates the transcription of autophagy markers, including Beclin-1, LC3b, and p62/SQSTM-1, thus upregulating the process of autophagy^16,63–65^. Previously, we demonstrated that NF-κB mediates the transcriptional upregulation of hnRNPD in oral cancer cells^34^. Interestingly, in the present study, we observed that the knockout of hnRNPD resulted in a drastic reduction in NF-κB expression in oral cancer cells, completely abolishing the expression of LC3b and p62/SQSTM-1. This led to the inhibition of autophagy and autophagic flux and the accumulation of autophagic vesicles in hnRNPD knockout oral cancer cells. These findings indicated that hnRNPD may be considered a new therapeutic target for the treatment of oral cancer. Our findings agree with previous studies, which suggest that the inhibition of autophagy in oral cancer could be used as targeted therapy for its treatment^66–70^. The PI3K/AKT/mTOR cell signaling pathway is a key regulator of autophagy. In the present study, we observed the constitutive activation of AKT that activated mTOR by phosphorylating it at the serine-2448 position. Phosphorylation of mTOR at this position is a marker for active PI3K/AKT pathway and mTOR kinase activity^71,72^. The active mTOR is an inhibitor of autophagy; however, in oral cancer cells, we observed constitutive activation of autophagy, which demonstrates that the activation of autophagy in oral cancer cells is not under the direct control of mTOR, but it is dependent on the NF-κB expression.

The hnRNPD KO resulted in the activation of cellular senescence observed by enhanced SA-β-Galactosidase staining in oral cancer cells. This finding is in support of a previous report in which hnRNPD KO mice showed premature aging through enhanced cellular senescence^22,36,73^.

### Conclusion

For the first time, the present study established the role of hnRNPD in the proliferation, migration, invasion, and colony formation of oral cancer. The present study’s findings demonstrated that autophagy is under the direct control of NF-κB in oral cancer cells. Our results indicated hnRNPD as a novel positive regulator of autophagy in oral cancer cells, and the hnRNPD KO resulted in the switching of cells from autophagy to cellular senescence.

## Abbreviations

HnRNPD: Heterogeneous Ribonucleoprotein D
AUF1: Adenylate Uridylate rich RNA binding Factor 1
PTEN: Phosphatase, and Tension homolog
PI3K: phosphatidylinositol-3 kinase
AKT: protein kinase B
ARE: Adenylate Uridylate Rich Element.

## Acknowledgment

The authors would like to acknowledge Dr. Stephanie J. Bouley, Dr. Priyanshu Sharma, and Dr. Abhimanyu Thakur for their help and critical insights in editing the final version of the manuscript.

## Author contributions

VK performed all experiments except immunohistochemistry and drafted the manuscript. AK and AT helped correct the manuscript. AM, PV, and VY helped in experiments. MRL performed immunohistochemistry. DP performed RNA-Seq data analysis. DM examined, reviewed, and scored immunohistochemistry. SSC and VK designed the study and wrote the final manuscript. SCC obtained funding.

## Funding

Department of Science and Technology (DST), Government of India, grant no. SERB/F/1230/2016-17, New Delhi, India, financially supported this study.

## Availability of data and materials

This published article includes all data generated or analysed during this study.

## Declarations

### Ethics Approval and Consent to Participate

Patient samples were collected after obtaining approval from the Institute Ethics Committee for Postgraduate Research, All India Institute of Medical Sciences, New Delhi, India (NO.IECPG-162/19.04.2018). Before sample collection, all participants and/or their legal guardian(s) gave written informed consent, a mandatory requirement from the institute’s ethics committee.

### Consent for publication

Not applicable.

### Competing interests

The authors declare they have no competing interests.

## Figure legends

**Supplementary Table 1:**
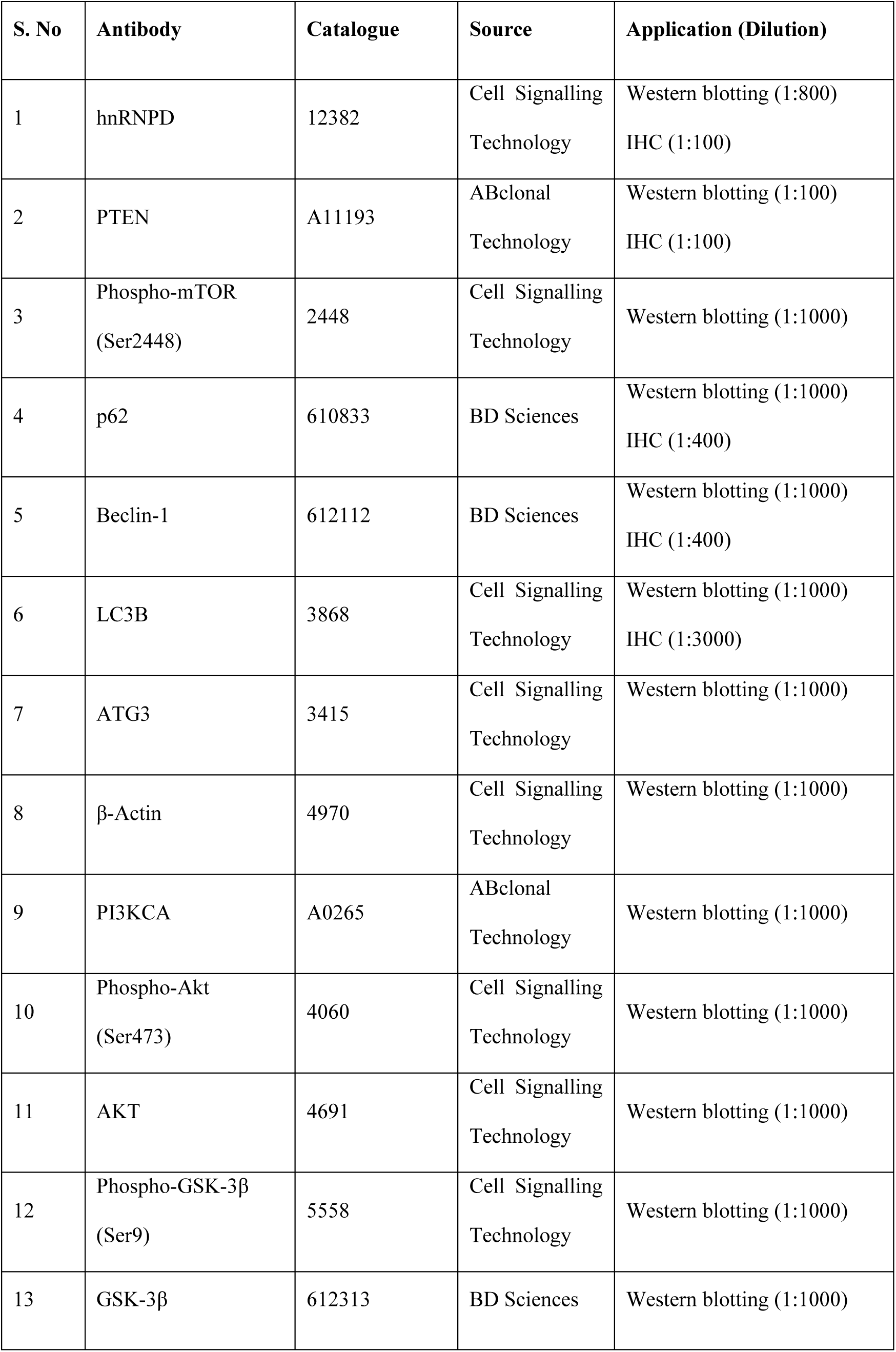
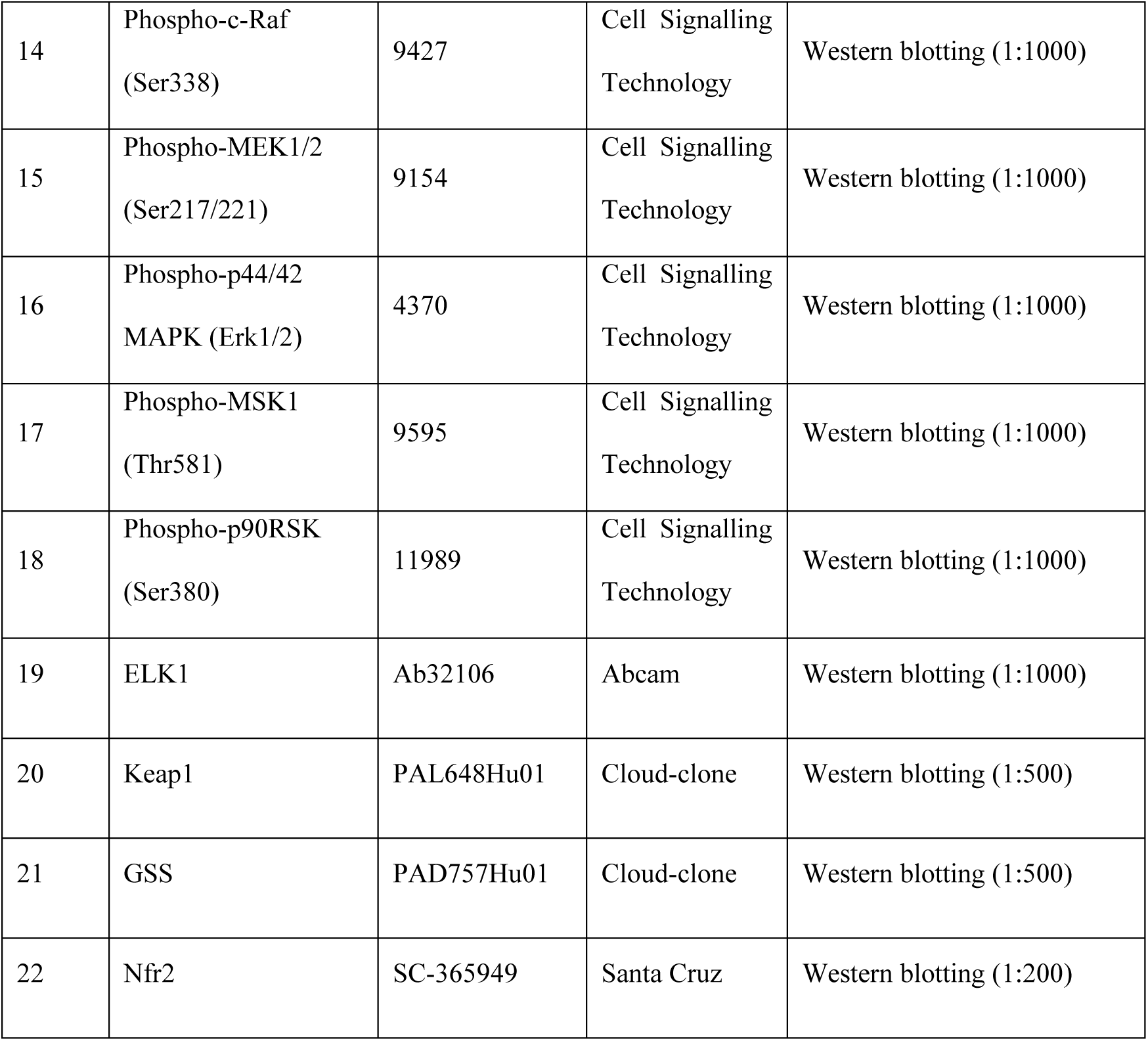
List of antibodies used in the present study.

**Supplementary Table 2:**
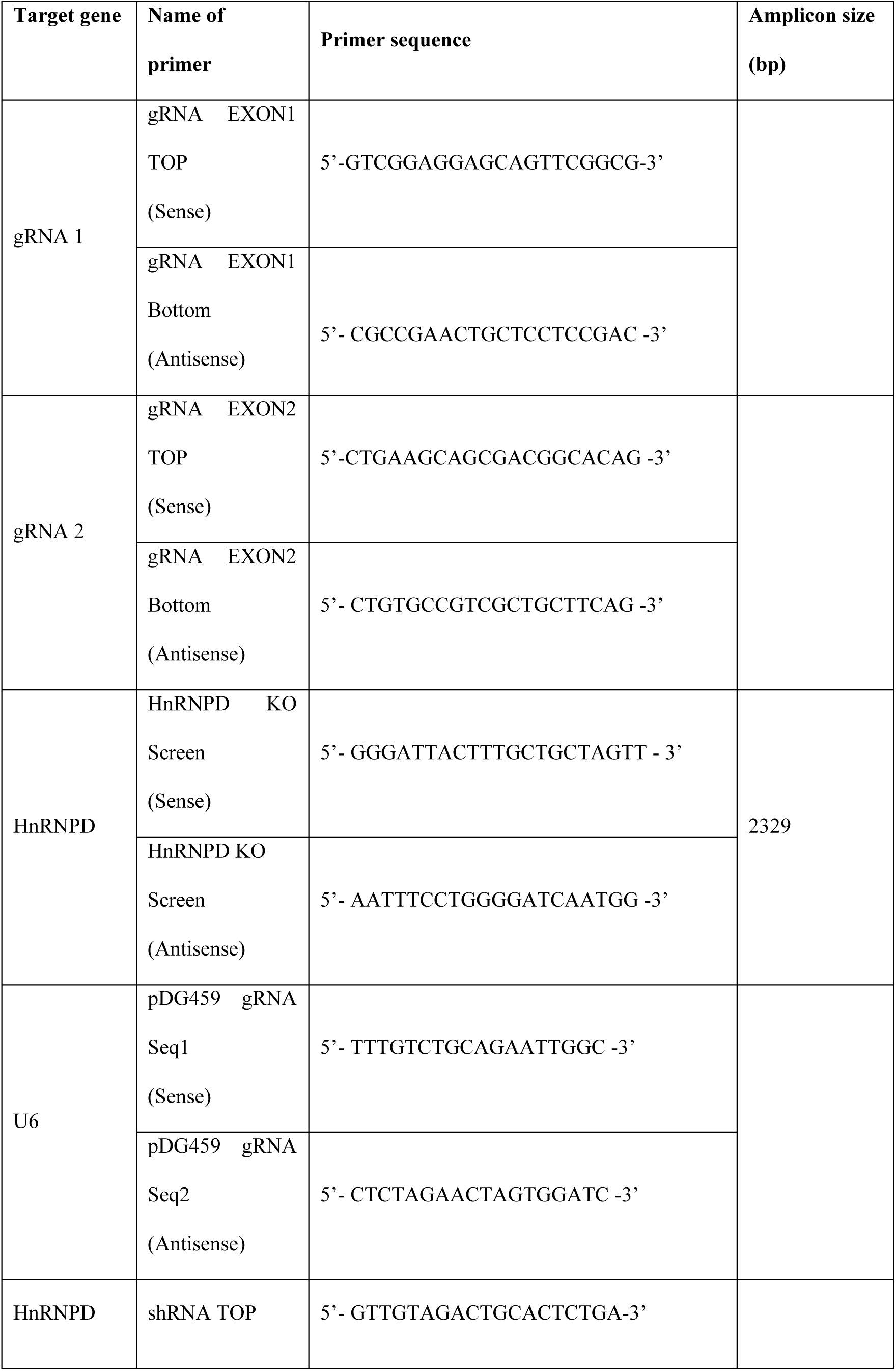
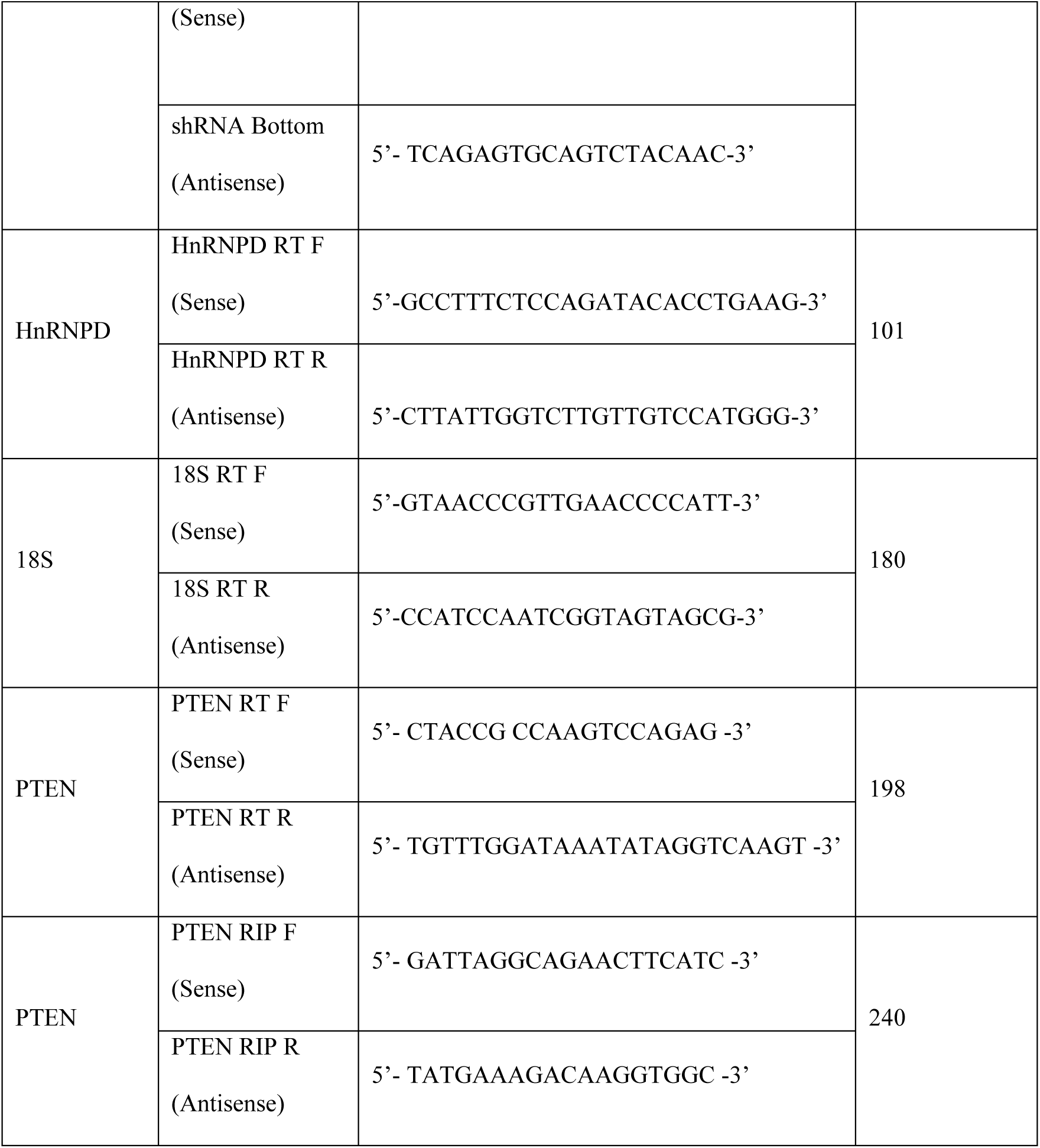
List of primers used in the present study.

**Supplementary Table 3:**
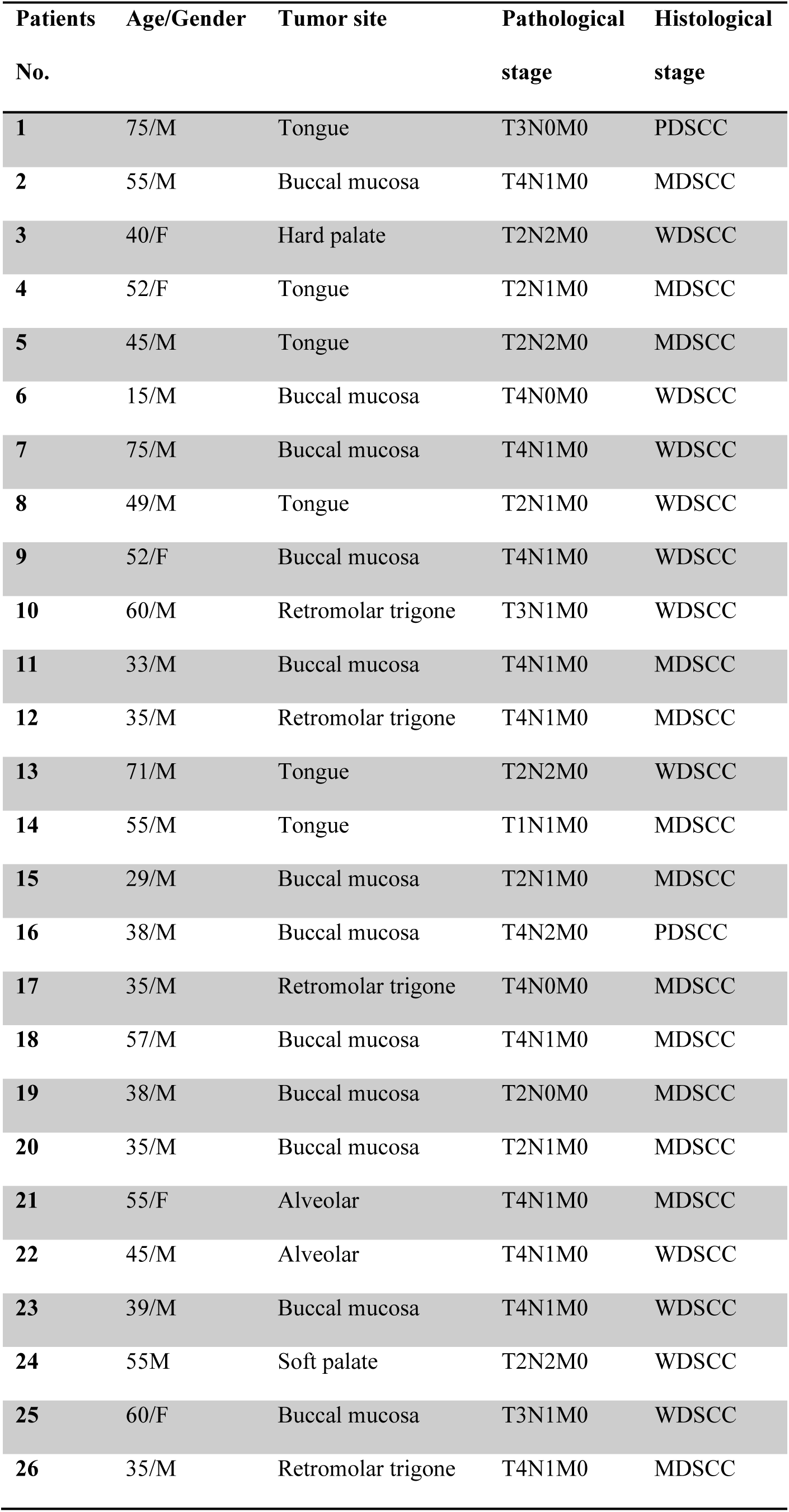
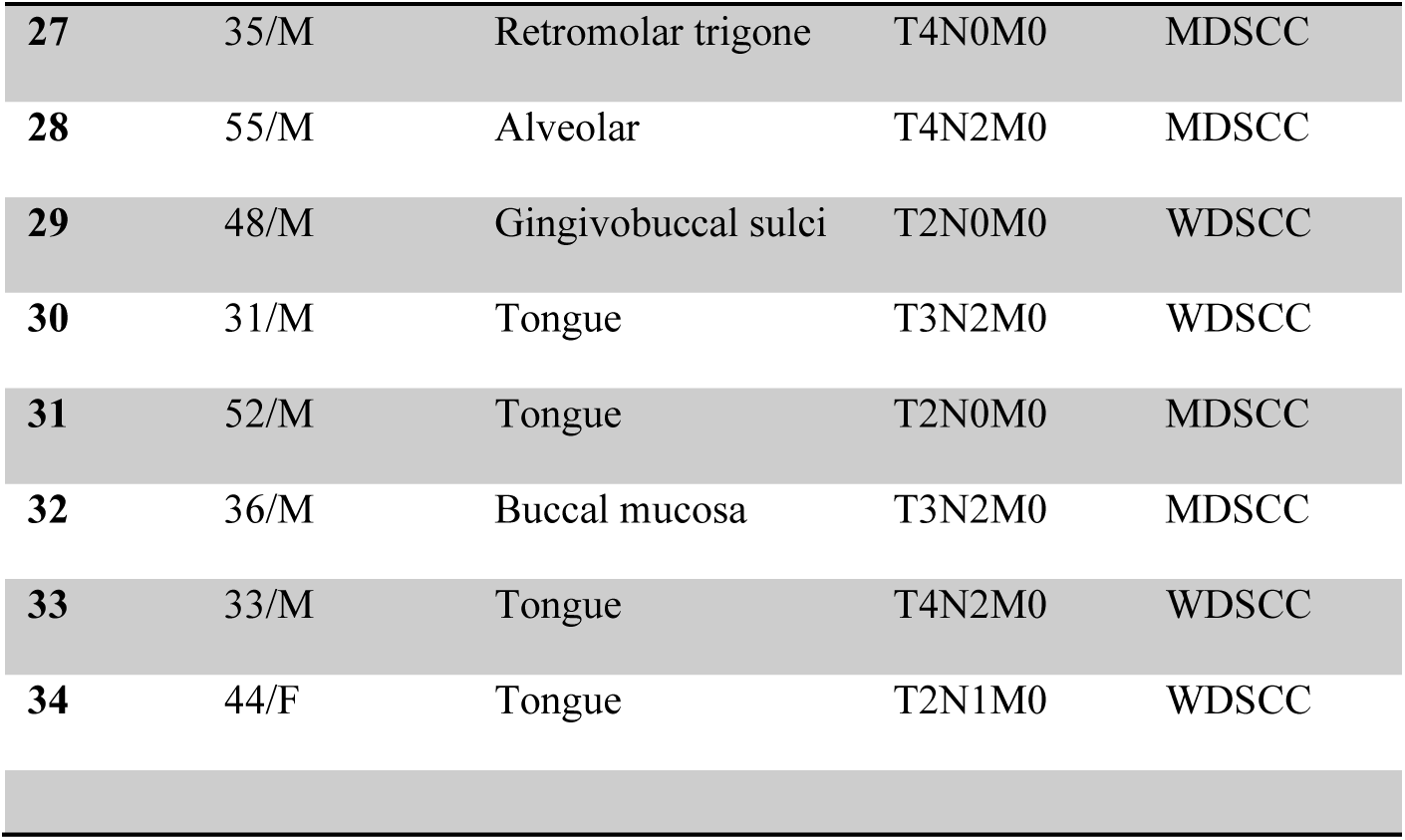
Clinical features of the patients.

**Figure S1:**
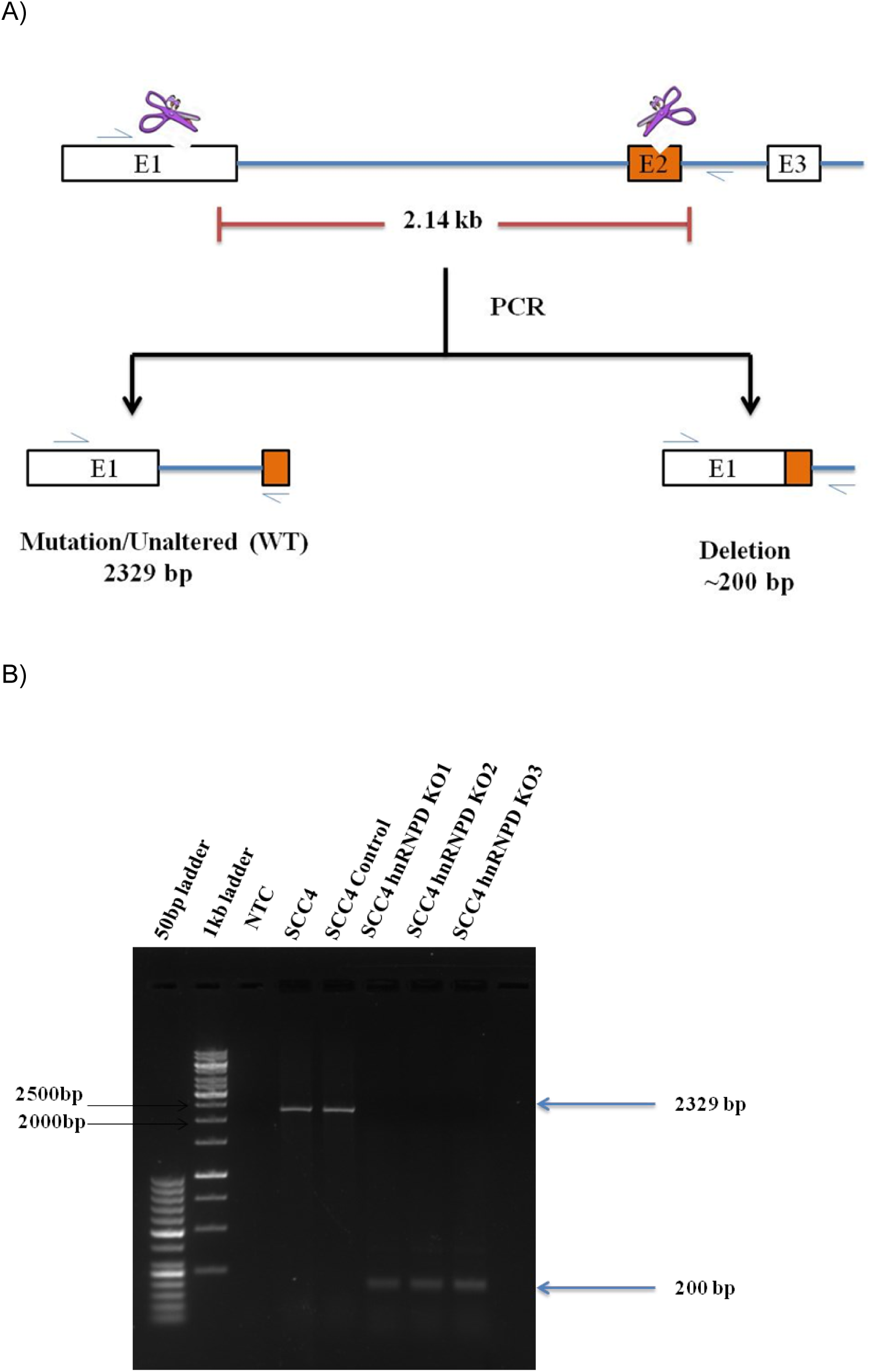
Strategy for CRISPR/Cas9 mediated knockout of human hnRNPD gene and identification of knockout clones by PCR-based screening. **(A)** The human hnRNPD gene on chromosome 4q21 consists of 9 exons and 8 introns. Alternative splicing of exons 2 and 7 leads to the synthesis of four different isoforms of hnRNPD^54^. Since all these isoforms perform overlapping functions, two guide RNA targeting exon1 and 2 were designed for knocking out all isoforms of hnRNPD gene-specific guide RNAs sequences, gRNA1 (Exon1; +316/+336), and gRNA2 (Exon2; +2434/+2454). These gRNAs were designed by using the CRISPick Genetic Perturbation Portal (https://portals.broadinstitute.org/gppx/crispick/public) an online available sgRNA Designer from the Broad Institute (MIT and Harvard, USA) and selected based on the highest cut target probability and lowest off-target rank. These gene-specific gRNAs sequences were cloned downstream to the individual U6 promoter, in the gRNA scaffolds by utilizing a one-step cloning protocol in the pDG459 plasmid. The nucleotide sequences of these gRNAs exhibited 100% homology with the 4q21 locus of the human genome on Exon1 (+316/+336), and Exon2 (+2434/+2454) of the hnRNPD gene. The resulting vector as transfected in oral squamous cancer cells SCC-4 followed by selection with 2µg/ml puromycin. The binding of the RNP complex (Cas9 nuclease and gRNAs) to the target regions resulted in the activation of cas9 nuclease, which mediates cleavage of DNA by introducing a double-stranded break in the target gene ∼3 bp upstream to the PAM sequence. Non-homologous end joining (NHEJ) repair pathway that is more error generous, this repair pathway introduces the addition or deletion of nucleotides. The antibiotic-resistant cells were collected and cloned by limiting dilution method in 96 well plates. Which were confirmed by PCR and DNA sequencing. The knockout of hnRNPD expression was further confirmed by western blotting. **(B)** Total genomic DNA isolated from hnRNPD knockout clones was isolated and subjected to PCR using hnRNPD KO screen sense and antisense which flanked guide RNA 1 and 2 respectively. The nucleotide sequences of these primers and their binding location on hnRNPD gene in table-2. The PCR product was resolved on 1.2% agarose and visualized after staining with ethidium bromide. Amplification of 2329 bp smaller DNA fragment in knockout cells compared to the control cells established the physical deletion of the hnRNPD gene between the target sites of two guide RNAs on exon1 and exon 2 respectively. Using this strategy, we successfully knocked out all the isoforms of hnRNPD in SCC-4 cells.

**Figure S2:**
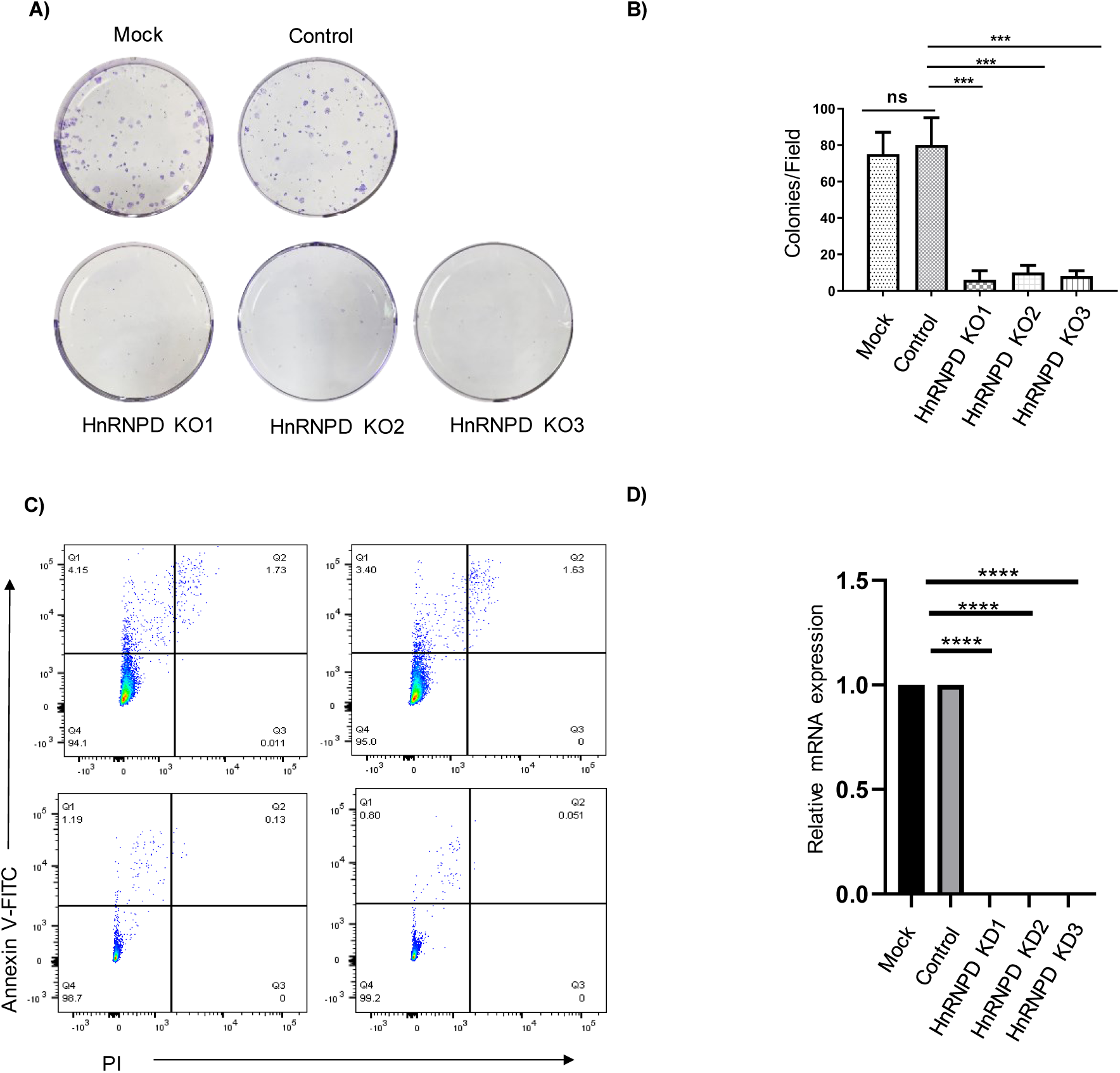
Knockout of hnRNPD leads to a reduction in oncogenic properties of oral cancer cells. **(A)** Representative photomicrographs of colony formation assay followed by crystal violet staining. **(B)** Number of colonies was counted in the four different fields and the mean±SD was plotted. **(C)** Scatterplots of cells stained with Annexin V/PI for apoptosis and necrosis. **(D)** Total RNA isolated from SCC-4 control and hnRNPD knockdown SCC-4 cells was reverse transcribed and subjected to real-time PCR using specific hnRNPD primers. Simultaneously, the PCR was also performed using 18S ribosomal RNA-specific primers and served as an internal control for normalization of the hnRNPD transcript.

## Notes

### Competing Interest Statement

The authors have declared no competing interest.

